# ORP5 AND ORP8 ORCHESTRATE LIPID DROPLET BIOGENESIS AND MAINTENANCE AT ER-MITOCHONDRIA CONTACT SITES

**DOI:** 10.1101/2021.11.11.468233

**Authors:** Valentin Guyard, Vera F. Monteiro-Cardoso, Mohyeddine Omrane, Cécile Sauvanet, Audrey Houcine, Claire Boulogne, Kalthoum Ben Mbarek, Nicolas Vitale, Orestis Facklaris, Naima El Khallouki, Abdou Rachid Thiam, Francesca Giordano

**Affiliations:** Institute for Integrative Biology of the Cell (I2BC), CEA, CNRS, Université Paris-Saclay, Gif-sur-Yvette cedex 91198, France; Inserm U1280, Gif-sur-Yvette cedex 91198, France; Laboratoire de Physique de l’École Normale Supérieure, ENS, Université PSL, CNRS, Sorbonne Université, Université de Paris, F-75005 Paris, France; Institut Jacques Monod, CNRS, UMR7592, Université Paris Diderot, Sorbonne Paris Cité, F-75013 Paris, France; Imagerie-Gif, Electron Microscopy Facility, Institute for Integrative Biology of the Cell (I2BC), CEA, CNRS, Univ. Paris-Sud, Université Paris-Saclay, Gif-sur-Yvette cedex 91198, France; Centre National de la Recherche Scientifique, Université de Strasbourg, Institut des Neurosciences Cellulaires et Intégratives, UPR-321267000 Strasbourg, France; MRI, BioCampus Montpellier, CRBM, Univ. Montpellier, CNRS, Montpellier, France

**Keywords:** Lipid droplets, organelle biogenesis, membrane contact sites, MAM, ORP, seipin, phosphatidic acid

## Abstract

Lipid droplets (LDs) are the primary organelles of lipid storage, buffering energy fluctuations of the cell. They store neutral lipids in their core that is surrounded by a protein-decorated phospholipid monolayer. LDs arise from the Endoplasmic Reticulum (ER). The ER-protein seipin, localizing at ER-LD junctions, controls LD nucleation and growth. However, how LD biogenesis is spatially and temporally coordinated remains elusive. Here, we show that the lipid transfer proteins ORP5 and ORP8 control LD biogenesis at Mitochondria-Associated ER Membrane (MAM) subdomains, enriched in phosphatidic acid. We found that ORP5/8 regulate seipin recruitment to these MAM-LD contacts, and their loss impairs LD biogenesis. Importantly, the integrity of ER-mitochondria contact sites is crucial for the ORP5/8 function in regulating seipin-mediated LD biogenesis. Our study uncovers an unprecedented ORP5/8 role in orchestrating LD biogenesis at MAMs and brings novel insights into the metabolic crosstalk between mitochondria, ER, and LDs at membrane contact sites.

**HIGHLIGHTS:**

ORP5 and ORP8 localize at MAM subdomains where LDs originate.
Phosphatidic acid is enriched in MAM subdomains that are the birthplace of LDs.
ORP5 and ORP8 knockdown impairs LD biogenesis.
ORP5 and ORP8 regulate seipin recruitment to MAM-LD contact sites.

## INTRODUCTION

Lipid Droplets (LDs) are evolutionarily conserved organelles that play a primary role in regulating lipid metabolism by storing lipids in excess and releasing them upon cellular needs (Olzmann and Carvalho, 2019). They store neutral lipids in their core, mainly triacylglycerol (TG) and sterol esters, surrounded by a protein-decorated phospholipid monolayer (Thiam et al., 2013). The storage of these lipids is essential for the cells to respond to energy fluctuations. Yet, LDs are also involved in other cellular functions such as protein degradation, gene expression regulation, lipid sequestration, and membrane biosynthesis (Welte and Gould, 2017).

LDs arise from the Endoplasmic Reticulum (ER) following the biosynthesis and deposition of neutral lipids in the bilayer hydrophobic region (Thiam and Ikonen, 2020). The neutral lipids condense to nucleate a nascent LD, which grows and buds into a mature LD. Defects in LD biogenesis and regulation are the hallmark of multiple metabolic and non-metabolic disorders such as type II diabetes, heart diseases, or viral infections (Gluchowski et al., 2017; Herker et al., 2021).

Seipin is a conserved ER protein forming a large oligomeric ring complex (Klug et al., 2021; Sui et al., 2018; Yan et al., 2018): it is a master regulator of LD biogenesis, and it plays a critical role in adipogenesis (Rao and Goodman, 2021). Mutations in seipin result in impaired lipid and calcium metabolism (Bi et al., 2014; Pagac et al., 2016) and cause lipodystrophies and neuronal disorders (Magré et al., 2001; Rao and Goodman, 2021; Windpassinger et al., 2004). Seipin physically marks the site of LD nucleation and mediates LD growth (Chung et al., 2019; Fei et al., 2008; Salo et al., 2019; Szymanski et al., 2007). It also regulates the physical connection of newly formed LDs to the ER, at ER-LD contact sites (Schuldiner and Bohnert, 2017), and facilitates LD growth (Choudhary et al., 2020; Grippa et al., 2015; Salo and Ikonen, 2019; Salo et al., 2019; Wang et al., 2016). In human cells, the oligomeric seipin ring consists of 11 subunits that bind negatively charged phospholipids, *in vitro*, including phosphatidic acid (PA) (Yan et al., 2018). In the absence of seipin, PA levels likely increase in the ER membrane (Fei et al., 2011; Han et al., 2015; Wolinski et al., 2015). Interestingly, seipin deletion leads to PA accumulation into the inner nuclear envelope and the formation of nuclear LDs (So\ltysik et al., 2021). These observations support the hypothesis that seipin is recruited to ER subdomains, probably enriched in negatively charged lipids (especially PA), where it could form a scaffold that assists LD assembly.

The existence of ER-subdomains promoting LD biogenesis is supported by several observations (Choudhary et al., 2020; Hariri et al., 2018; Joshi et al., 2018; Santinho et al., 2020; Wang et al., 2018). LD biogenesis can occur at the ER contact sites with vacuoles in yeast (Hariri et al., 2018) or peroxisomes in yeast and mammals (Joshi et al., 2018, 2021). Based on these observations, contact sites established between the ER and other organelles could play a critical role in LD formation. For example, such contact sites may pool key enzymes and lipid intermediates necessary for LD assembly (Choudhary et al., 2020) or preset optimal local physical properties for the neutral lipids’ condensation into LDs. Whether LD can originate at other inter-organelle contact sites is unknown.

Perturbations of the lipid composition in ER are detrimental to proper LD formation (Adeyo et al., 2011; Ben M’barek et al., 2017; Fei et al., 2011; Zoni et al., 2021). Therefore, locally editing the ER phospholipid composition, possibly by lipid transfer at ER-LD contact sites, could be essential for LD biogenesis. However, the existence of such a mechanism is currently unknown. Also, it is still elusive whether and how lipids are directly transferred from the ER to the LDs during and after LD formation. Therefore, mechanisms that spatially and temporally regulate LD biogenesis in the cell remain poorly understood.

The Oxysterol binding protein (OSBP)-related proteins constitute a large family of lipid transfer proteins (LTPs) conserved from yeast (Osh) to humans (ORP) and localized to different subcellular sites, shown in several cases to be membrane contact sites. A common feature of all ORPs is the presence of an OSBP-related lipid-binding/transfer (ORD) domain. Most ORP proteins contain a two phenylalanines (FF) in an acidic tract (FFAT)-motif that binds ER-localized VAP proteins and a pleckstrin homology (PH) domain that interacts with lipids or proteins in distinct non-ER organelle membranes. Two members of this family, ORP5 and ORP8, do not contain an FFAT motif but are directly anchored to the ER through a C-terminal transmembrane (TM) segment.

ORP5 and ORP8 have been originally shown to localize at ER-PM contact sites where they transfer phosphatidylserine (PS) from the cortical ER to the PM, in counter-exchange with the phosphoinositides Phosphatidylinositol-4-phosphate (PtdIns(4)P) and Phosphatidylinositol 4,5-bisphosphate (PtdIns(4,5)P) (Chung et al., 2015; Ghai et al., 2017). Lately, ORP5 has been shown to also localize at ER-LD contact sites (Du et al., 2020) and, although an experimental demonstration is still missing, it has been proposed that, at these sites, ORP5 could also act as PS/PtdIns(4)P lipid exchanger.

We recently showed that ORP5 and ORP8 are also localized, and even enriched, at a specific ER subdomain in contact with mitochondria, the so-called mitochondria-associated membranes (MAM), where they mediate transfer of PS, maintain mitochondrial morphology and function, and control calcium fluxes (Galmes et al., 2016; Rochin et al., 2019). We hypothesized that MAMs containing ORP5 and ORP8 are involved in LD biogenesis.

Here, we show that ORP5 and ORP8 localize at MAM subdomains where LDs originate and that are enriched in PA lipid. Oleic acid treatment leads to massive recruitment of ORP5-labeled MAM to nascent and pre-existent LDs, suggesting a dual role in LD biogenesis and maintenance/turnover. We also reveal that ORP5 and ORP8 are novel players involved in LD biogenesis by regulating seipin recruitment to MAM-LD contacts.

## RESULTS

### ORP5 localize at MAM subdomains closely associated with LDs

ORP5 and ORP8 were originally identified as components of ER-PM contact sites (Chung et al., 2015). They have been recently shown to also localize at contact sites between ER and mitochondria (Mito) (Galmes et al., 2016) and to primarily interact at MAMs (Rochin et al., 2021). Lately, a novel localization at ER-LD contact sites upon oleic acid (OA) treatment has been described for the two ORP5 isoforms (ORP5A and ORP5B) but not for ORP8 (Du et al., 2020). However, ORP5 distribution at the multiple contact sites, especially upon OA, remains controversial. Also, whether ORP8 could also localize at ER-LDs contact sites is unknown.

Thus, we first investigated the localization of EGFP-tagged ORP5A, ORP5B (a natural variant of ORP5, lacking a large part of the PH domain), and ORP5ΔPH (a variant completely lacking its PH domain) by confocal microscopy in HeLa cells treated for 2h with OA or untreated. LipidTox (LTox) was used to stain LDs and mitotracker to label mitochondria (Fig 1A-B, S1A).

**Figure 1.**
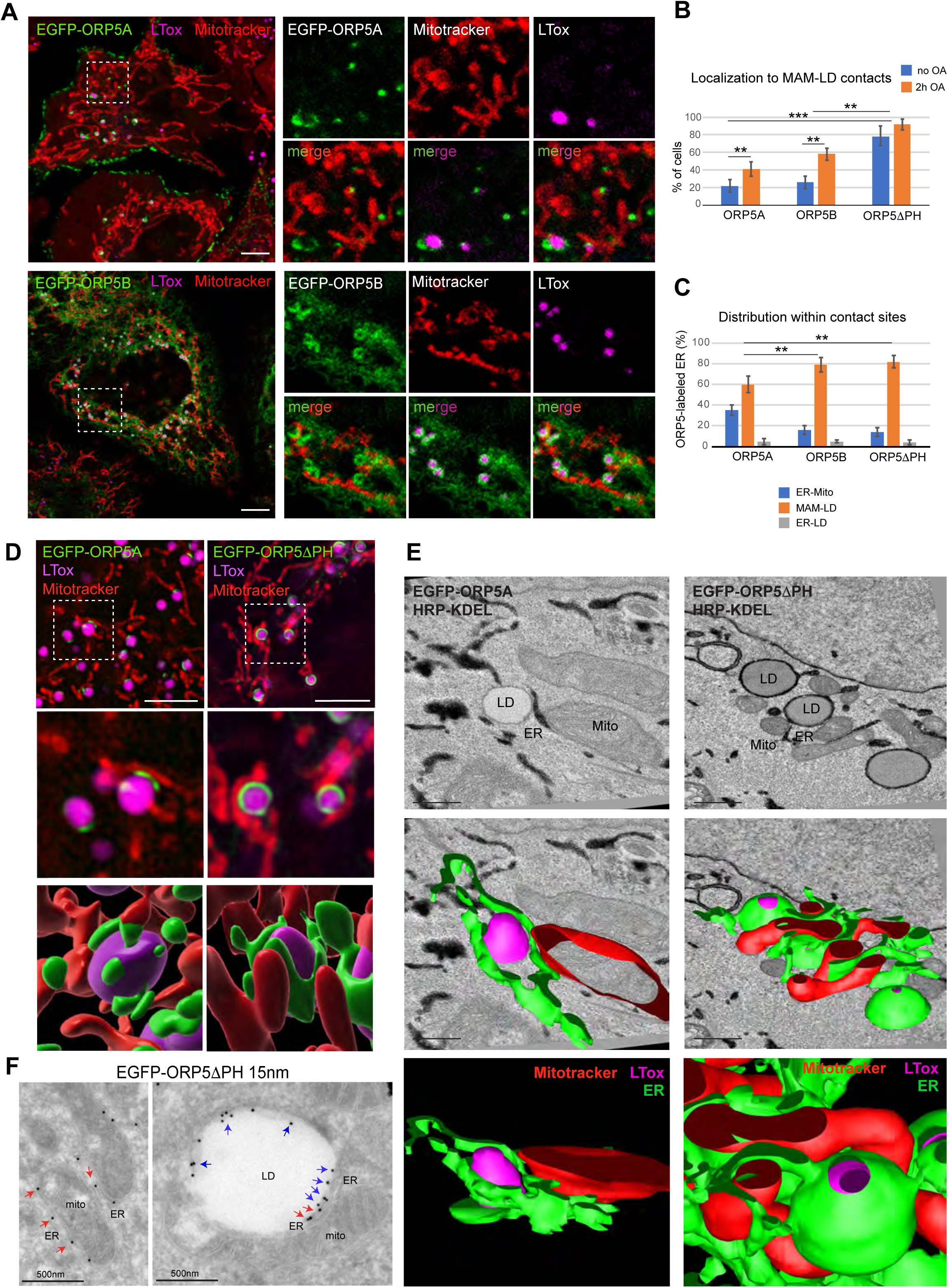
ORP5 localizes to MAM subdomains in contact with LD. **A)** Representative confocal images showing single focal planes of HeLa cells expressing EGFP-ORP5A or EGFP-ORP5B (green) and treated with Mitotracker (red) and LTox Deep Red (LTox, purple) to label mitochondria and lipid droplets (LD), respectively. Scale bar 10 μm. **B)** Quantification of the % of HeLa cells showing localization of EGFP-tagged ORP5A, ORP5B and ORP5ΔPH to MAM-LD contacts, in absence of Oleic acid (OA) treatment or after 2hr of 300 μM OA treatment. Data represent the mean±standard error of the mean (SEM) of n= 25 cells. ** p<0.01, ***p<0.001, unpaired two-tailed *t*-test. **C)** Analysis of the distribution of EGFP-tagged ORP5A, ORP5B and ORP5ΔPH among the contact sites ER (MAM)-mitochondria (Mito), MAM-LD and ER-LD in HeLa cells treated with 300 μM OA for 2hr. Data represent the mean±SEM of n=15 cells. ** p<0.01. **D)** Structured illumination microscopy (SIM) micrographs of HeLa cells expressing EGFP-ORP5B or EGFP-ORP5ΔPH (green), treated with OA for 2hr, and stained with Mitotracker (red) and LTox Deep Red (purple). 3D-SIM images were obtained by segmentation using software Imaris (v 9.3, Bitplaine). Scale bar, 5μm. **E)** Representative electron micrographs of HeLa cells co-overexpressing EGFP-ORP5A or EGFP-ORP5ΔPH and HRP-KDEL after 2hr OA treatment, and their 3D representation obtained by 3Dmod. LD, lipid droplet, Mito, mitochondria, and ER, endoplasmic reticulum. Scale bar 500nm. **F)** Electron micrograph of ultrathin cryosections of HeLa cells transfected with EGFP-ORP5ΔPH, treated with OA for 2hr, and immunogold stained with anti-EGFP (15 nm gold). Left: red arrows indicate EGFP-ORP5ΔPH localized to MAM. Right: blue arrows indicate ORP5ΔPH localized to ER-LD contacts and red arrows indicate ORP5ΔPH localized to MAM-LD contacts. Scale bar 500 nm.

EGFP-ORP5A was detected at cortical ER (ER-PM contacts) in all cells analyzed (OA treated and untreated), while only a subset of cells displayed ORP5A localization at ER-LD contacts (20% for untreated and 40% for OA-treated cells). Interestingly, the ORP5 labeled-ER in contact with LDs appeared to be highly expanded as it almost completely surrounded the LD surface. EGFP-ORP5B was instead detected at reticular ER (due to the loss of PM binding) in all cells analyzed and at expanded ER-LD contact sites in 22% of untreated cells and in about 60% of OA-treated cells. Similarly, ORP5ΔPH was found at ER-LDs contact sites. However, the number of cells showing ORP5-labeled ER in contact with LD was considerably higher in the pool of cells transfected with ORP5ΔPH (80% for untreated and 96% for OA-treated cells) than in the pool of cells transfected with EGFP-ORP5B (Fig 1B and S1A).

Interestingly, almost all the ER subdomains in contact with LDs where ORP5A and ORP5B localized were found closely associated to mitochondria, forming a novel three-way Mito-MAM-LD membrane contact site structure (Fig 1A). Quantification of these observations revealed that 60% of EGFP-ORP5A-, 80% of EGFP-ORP5B- and 80% EGFP-ORP5ΔPH-positive ER structures corresponded to Mito-MAM-LD contact sites, while 35% of EGFP-ORP5A-, 15% of EGFP-ORP5B- and 15% EGFP- ORP5ΔPH-positive ER structures corresponded to MAM. Only a small pool of EGFP- ORP5A-, of EGFP-ORP5B- and EGFP-ORP5ΔPH-positive ER structures (4%, 4%, and 3%, respectively) were associated only to LDs (Fig 1C).

To better visualize the three organelles’ association, we proceeded by cell swelling. Cells were subjected to OA for 2hr and then to a hypotonic medium, which swells the bilayer-surrounded organelles only. We found that 70% of LDs formed were in close association with mitochondria and ER vesicles; the remaining 30% were in contact with ER only (Fig S1B). This observation confirmed the development of Mito-MAM-LD contact sites upon OA addition. Interestingly, the pool of ORP5 at the MAM-LD contact was 15 times higher than in the ER regions, indicating that OA addition massively relocalizes ORP5B from the ER to MAM subdomains in contact with LDs.

Furthermore, super-resolution Structured Illuminated Microscopy (SIM) analysis of EGFP-ORP5A and EGFP-ORP5ΔPH localization in HeLa cells treated for 2h with OA confirmed their localization at Mito-MAM-LD close associations (Fig 1D).

To further study these three-way Mito-MAM-LD associations, we assessed their morphology through ultrastructural analysis by HRP-KDEL EM (carrying a horseradish peroxidase (HRP) tagged with an ER retention motif to stain the ER) in cells expressing HRP-KDEL alone or together with EGFP-ORP5A or EGFP-ORP5ΔPH (Fig 1E, Fig S1C). Our morphological analysis confirmed that the Mito-MAM-LD associations observed by light microscopy were indeed membrane contact sites (in the range of 10-30nm). Agreeing with our confocal data, ORP5A, and ORP5ΔPH to a greater extent, induced an expansion of the ER on the LD surface. Mito-MAM-LD associations were also detected in cells expressing HRP-KDEL alone, revealing the existence of this structure even in the absence of ORP5 overexpression. However, the ER involved in these contact sites was not highly-expanded around the LDs (Fig 1E, Fig S1C), possibly explaining why these associations have been overlooked so far.

Moreover, the localization of ORP5 at the expanded Mito-MAM-LD contact sites was confirmed by immunogold labeling on ultrathin cryosections of cells expressing ORP5ΔPH and treated with OA. The advantage of using ORP5ΔPH over ORP5A or ORP5B relies on the higher number of cells showing its localization at Mito-MAM-LD contact sites (see Fig 1B), facilitating the immuno-EM analysis (Fig 1F). The bulk of ORP5ΔPH localized to Mito-MAM-LD contact sites and to the associated ER-LD contacts, while a minor pool was detected at MAM (ER-mitochondria contacts) not associated with LDs (Fig 1F).

Together, these data reveal the existence of a novel tripartite Mito-MAM-LD contact sites junction where ORP5 localizes. However, it is not known whether its binding partner ORP8 also localizes at Mito-MAM-LD contact sites.

### ORP8 is enrichment at MAM-LD via interactions with ORP5 by its coiled-coil domain

We overexpressed EGFP-ORP8 and analyzed its localization by confocal microscopy in HeLa cells treated with OA (Fig 2A). In contrast to ORP5, ORP8 mostly localized to the reticular ER, with a minor pool additionally present at cortical ER and MAMs, as previously shown (Galmes et al., 2016; Rochin L, 2020). A small pool of ORP8 was detected at MAM-LD contact sites, but the ER elements labeled with ORP8 were not expanded around the LD surface as seen for ORP5.

**Figure 2.**
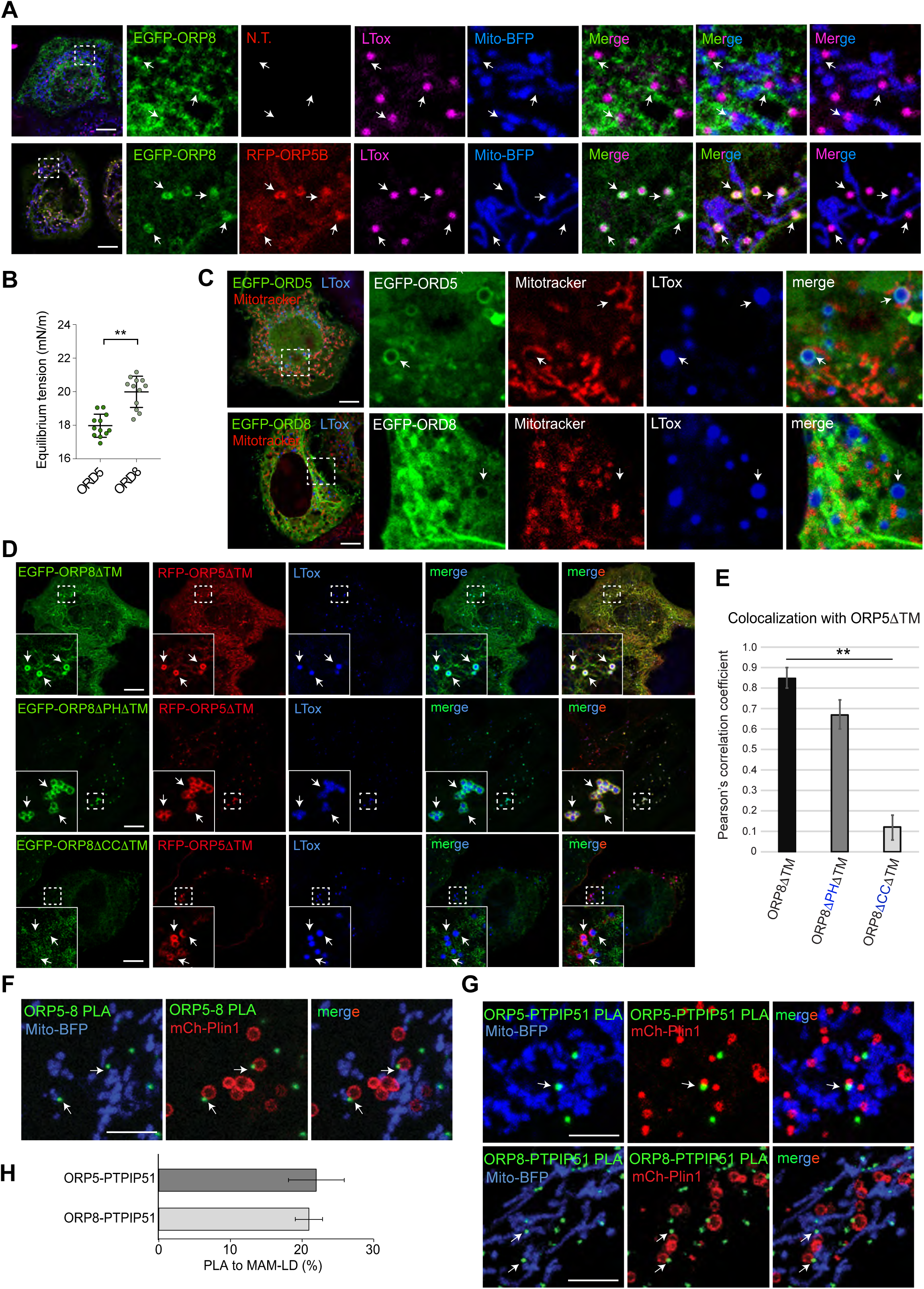
ORP8 localizes and interacts with ORP5 at Mito-MAM-LD contacts. **A)** Representative confocal images showing localization of EGFP-ORP8 alone (green) or together with RFP-ORP5B (red), in HeLa cells co-expressing Mito-BFP (blue), and treated with OA for 2hr and stained with LTox Deep Red (purple). Each image represents a single focal plane of confocal 3D stacks. Arrows point to ORP5-labeled MAM associated to mitochondria and LD (MAM-LD contacts). Scale bar 10 μm. **B)** LD- binding affinity of ORP5 and ORP8 ORD domains (ORD5 and ORD8, respectively) analyzed by tensiometry. Data are shown as mean±SEM of n=12, with ** p<0.01, unpaired Kolmogorov-Smirnov test. **C)** Confocal images showing the localization of EGFP-ORD5 and EGFP-ORD8 (green) in HeLa cells treated with OA for 2hr and stained with Mitotracker (red) and LTox Deep Red (blue). Each image represents a single focal plane, scale bar 10 μm. **D)** HeLa cells co-expressing EGFP-ORP8ΔTM and RFP-ORP5ΔTM, EGFP-ORP8ΔPHΔTM and RFP-ORP5ΔTM, or EGFP-ORP8ΔCCΔTM and RFP-ORP5ΔTM. Each image represents a single focal plane of confocal 3D stacks. Arrows point to ORP5 localization on the LD surface. Scale bar 10 μm. **E)** Quantitative analysis of the co-localization of either EGFP-ORP8ΔTM, EGFP-ORP8ΔPHΔTM or EGFP-ORP8ΔCCΔTM with RFP-ORP5ΔTM by Pearson’s correlation coefficient. Data represent mean±SEM of n=10 cells. **p<0.001 unpaired student’s *t*-test. **F-G)** Confocal micrographs showing endogenous ORP5-ORP8, ORP5-PTPIP51 and ORP8-PTPIP51 PLA interactions (green dots) in regions of HeLa cells co-expressing Mito-BFP (blue) and mCherry-Plin1 (mCh-Plin1) and treated with 300 μM OA for 2hr. Arrows point to PLA dots associated to MAM-LD contacts. Images represents a single focal plane. Scale bar 10 μm. **H)** Quantification of endogenous ORP5-PTPIP51 and ORP8-PTPIP51 PLA interaction at MAM-LD contacts. Data is shown as % mean±SEM of n= 36 cells (ORP5-PTPIP51) and n=15 cells (ORP8-PTPIP51).

ORP5 and ORP8 interact at MAMs (Rochin et al., 2021). We thus reasoned that the faint signal of ORP8 detected at the MAM-LD contact in overexpression conditions could be due to a lack of ORP5. We then overexpressed EGFP-ORP8 with RFP-ORP5B. Under this condition, ORP8 dramatically redistributes from the reticular and cortical ER to the ORP5-positive MAM-LD contact sites. This redistribution and enrichment at MAM-LD contacts were not observed in the case of a general ER protein such as Sec61b, which, even when co-expressed with ORP5, maintained its localization to the reticular ER (Fig S2A). These findings confirmed the specificity of ORP8 localization at MAM-LDs. Altogether, these data indicate that ORP5 has a higher affinity for MAM-LD contacts than ORP8, and that ORP8 enrichment at such sites depends on its interaction with and on the expression level of ORP5.

Deletion of the ORD domain from ORP5 eliminates its ability to localize at MAM-LDs (Fig S2B). This is in accord with recent data showing that the ORD domain of ORP5 is required for its localization to ER-LDs contacts (Du et al., 2020). How and whether this domain in ORP5/8 directly bind to LD surfaces is unknown. Thus, we decided to test *in vitro* whether these cytosolic ORD domains can interact with the neutral lipid/water interface of LDs and to compare their binding affinities. We used a tensiometer approach where we generated a triolein-in-buffer droplet (Ajjaji et al., 2019). The tension of this interface was 34mN/m, and then we added either the ORD domain of ORP5 or ORP8 at the same concentration. Adsorption of these domains to the interface will decrease tension; the better the adsorption, the higher the decrease in tension. We found that the ORD domain of ORP5 systematically decreased much better tension, from 34 to 18mN/m, than of ORP8, from 34mN/m to 20mN/m (Fig 2B). These experiments revealed that both ORD5 and ORD8 could bind LDs, but the binding ability of ORD8 would be lower than ORD5. Expression of EGFP-tagged ORDs in HeLa cells confirmed that the ORD domain of ORP5 (EGFP-ORD5) strongly binds LDs, while the ORD domain of ORP8 (EGFP-ORD8) only weakly binds LDs (arrows, Fig 2C). These data might explain why ORP8 overexpression does not induce an expansion of ER around the LDs and also suggest that ORP8 requires ORP5 to be enriched at the expanded MAM-LDs contact sites.

To more directly test whether ORP8 binding to LDs requires ORP5 and the domains involved, we co-expressed an ORP5 deletion mutant lacking the ER-anchoring TM domain (RFP-ORP5ΔTM) with an ORP8 construct carrying a similar mutation (EGFP-ORP8ΔTM) or with ORP8 lacking both the TM and the PH or the coiled-coil (CC) domains (EGFP-ORP8-ΔPHΔTM or EGFP-ORP8-ΔCCΔTM) and analyzed their recruitment to LDs in HeLa cells by confocal microscopy (Fig 2D). The EGFP-ORP5ΔTM, when expressed alone, localized to both the PM and LDs, while the EGFP-ORP8ΔTM mostly localized to the cytosol, to the PM, and weakly to LDs (Fig S2C). However, when co-expressed with RFP-ORP5ΔTM, EGFP-ORP8ΔTM strongly co-localizes with ORP5, as assessed by the Pearson’s correlation coefficient analysis, and enriches at the ORP5-labeled LDs (Fig 2D-E). Of note, ORP5ΔTM and ORP8ΔTM, when co-expressed, lose their PM localization and strongly redistribute to LDs, and also to cytosolic filamentous structures that resemble cytoskeleton.

The concomitant deletion of the PH domain in ORP8ΔTM does not induce significant changes in ORP8 colocalization with ORP5 at LDs. However, the deletion of the CC domain in ORP8ΔTM completely abolished ORP8 targeting to the ORP5-labeled LDs (Fig 2D-E). These results confirm that interaction with ORP5 via the CC domain is required for the ORP8 binding to LDs.

Finally, we determined whether ORP5 and ORP8 interaction and localization at MAM-LD contacts also occur at endogenous levels. We performed Proximity Ligation Assay (PLA) by duolink in HeLa cells transfected with Mito-BFP, to stain the mitochondria and with mCherry-Perilipin 1 (Plin1), to stain the LDs, and analyzed by confocal microscopy. ORP5-ORP8 PLA spots were detected in the proximity of a subset of mitochondria-associated LDs (arrows, Fig 2F). To confirm that these sites corresponded to MAMs, we performed PLA using antibodies against either ORP5 or ORP8 and their mitochondrial binding partner PTPIP51 (Galmes et al., 2016). A similar pool of ORP8-PTPIP51 or ORP5-PTPIP51 PLA spots was observed at MAM-LD contacts (arrows, Fig 2G-H), confirming that ORP5-8 localize and interact at the tripartite Mito-MAM-LD contact sites in endogenous conditions.

### ORP5 and ORP8 organize LD biogenesis at MAM

To assess the role of ORP5 and ORP8 in LD biogenesis, we depleted ORP5 and ORP8 by RNAi in HeLa cells that have been delipidated for 3 days to remove pre-existent LDs. The efficiency of the knockdown (KD) was validated by western blot (WB) (Fig S3A-B). LD biogenesis was induced by OA treatment, and cells were imaged by confocal microscopy at different times (15min, 30min, 1hr, 2hr) (Fig S3A). The abundance of LDs, stained by LTox, in both ORP5 and ORP8 KD cells was significantly reduced at all time points, as compared with control (Ctrl) cells. The decrease in LD number was greater at earlier times (70% at 15 min, 60-65% at 30 min and 1hr, 30-40% at 2hr), suggesting a delay in LD biogenesis (Fig S3A, S3C). Since LTox may have marked pre-existing LDs that could have resisted delipidation, we performed parallel experiments by feeding cells micelles containing a fluorescent C_12_-fatty acid (referred to as FA^568^, associated with red fluorescence), as in (Khaldoun et al., 2014). This option enabled us to track efficiently the biogenesis and the maturation of newly synthesized LDs. The efficiency of delipidation was confirmed by the almost complete disappearance of LTox-positive LDs in Ctrl, ORP5, and ORP8 KD cells at time=0 min (Fig 3A, 3C). The FA^568^ treatment induced the formation of new LDs that were also labeled by LTox. The number of the FA^568^–containing LDs was dramatically reduced at all time points (86% at 15 min, 92% at 30 min, and 71% at 1hr) in ORP5 and ORP8 KD cells as compared with control cells. The decrease in newly-formed FA^568^–positive LDs was higher than the decrease in LTox-positive LDs, which likely included the pre-existent ones (Fig 3B-C). These results indicate a role of ORP5 and ORP8 in regulating LD biogenesis.

**Figure 3.**
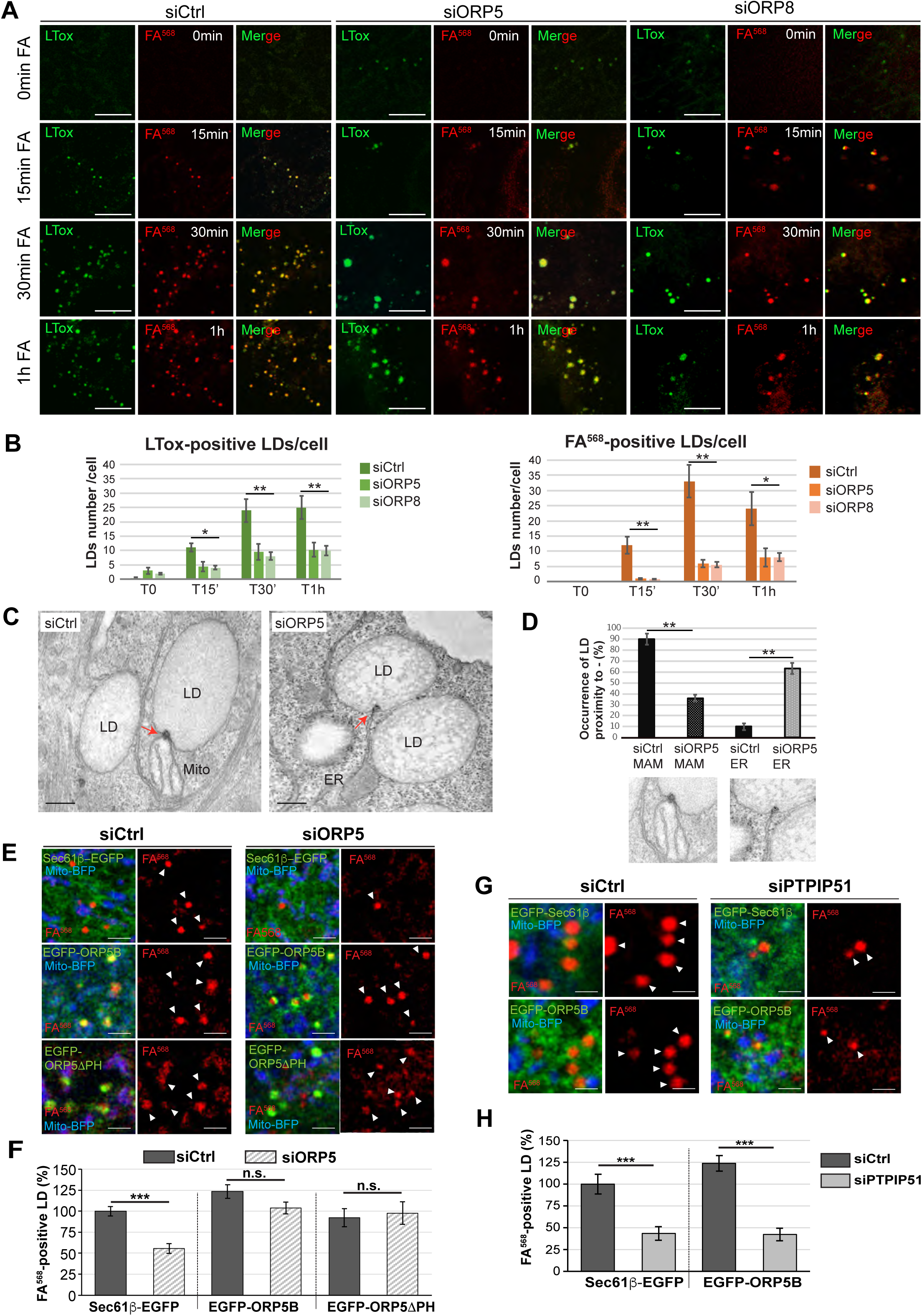
Depletion of ORP5 and ORP8 affect LD biogenesis. **A)** LD biogenesis time-course. HeLa cells delipidated for 72hr were treated with siCtrl, siORP5 or siORP8, incubated with 1μM FA^568^ (red) and stained with LTox Deep Red (green). Representative confocal images of regions of HeLa cells submitted to these experimental conditions at time 0 min, 15 min, 30 min and 1 hr of FA^568^ incubation are displayed as single focal plane. Scale bar 5 μm. **B)** Quantification of the number of FA^568^- and LTox-positive LD in control, ORP5 and ORP8 knockdown HeLa cells at the indicated times. Data represent mean±SEM of n= 30 cells. *p<0.001, **p<0.0001, unpaired two-tailed *t*-test. **C)** Representative electron micrographs of control and ORP5 knockdown HeLa cells, evidencing (red arrows) the electrondense structure that connects the nascent LD to the ER from which it originated and sometimes also to the mitochondria (Mito) at MAM-LD contacts. Scale bar 250 nm. **D)** Quantification of the number of LD associated to these ER or MAM electrondense structures. Data represent mean±SEM of n= 20 cells. **p<0.0001, unpaired two-tailed *t*-test. **E)** Confocal (single focal plane) micrographs of regions of control and ORP5 knockdown delipidated HeLa cells co-overexpressing Mito-BFP (blue) with either Sec61β-EGFP (green), or EGFP-ORP5B (green), or EGFP-ORP5ΔPH (green). Arrowheads indicate the newly formed LD. Scale bar, 2 μm. **F)** Quantitative analysis of the number of FA^568^- positive LD in control and ORP5 knockdown delipidated HeLa cells co-overexpressing Mito-BFP and either Sec61β-EGFP, or siRNA-resistants EGFP-ORP5B or EGFP- ORP5ΔPH, and treated for 15 min with FA^568^. Data are shown as % of mean±SEM of n=20-85 cells. ***p<0.001, unpaired two-tailed *t*-test. **G)** Confocal (single focal plane) micrographs of regions of control and PTPIP51 knockdown delipidated Hela cells, co-overexpressing Mito-BFP (blue) with Sec61β-EGFP (green) or EGFP-ORP5B (green) and treated for 1hr with FA^568^. Arrowheads indicate the newly formed LD. Scale bar, 1 μm. **F)** Quantification of the number of FA^568^-positive LD in control and PTPIP51 knockdown delipidated HeLa cell co-overexpressing Mito-BFP and either Sec61β- EGFP or EGFP-ORP5B and treated for 1hr with FA^568^. Data are shown as % of mean±SEM of n=20-22 cells. ***p<0.001, unpaired two-tailed *t*-test.

To analyze the morphology of nascent LDs at MAMs, we performed EM analysis on control and ORP5 KD cells treated with OA. Importantly, we detected specific ER subdomains that appeared as peculiar electrondense structures associated with some LDs connected to tubular ER elements, and that likely correspond to the sites where these LD emerged from the ER. These ER subdomains very often corresponded to MAM in control cells (Fig 3C). Quantifications of these EM observations revealed that ORP5 KD induces a strong decrease in the occurrence of these MAM-emerged LDs while the ER-emerged LD connections, not in close contact with mitochondria, were instead increased (Fig 3D). These data strongly suggest that ORP5 regulates LD formation from the MAM subdomain.

We then tested if re-expression of ORP5 could rescue the LD phenotype of the ORP5 KD cells. To this purpose, we re-expressed siRNA-resistant EGFP-ORP5B or EGFP-ORP5ΔPH or, as a control, the ER marker protein EGFP-Sec61β in the delipidated ORP5 KD cells. To monitor rescue of the LD phenotype, we performed FA^568^-mediated induction of LD biogenesis, as described above, and analyzed by confocal microscopy the number of FA^568^–positive LDs at 15 minutes after its delivery. Remarkably, both EGFP-ORP5B and EGFP-ORP5ΔPH constructs significantly rescued the LD decrease in siORP5 cells, while EGFP-Sec61β was not able to rescue the LD phenotype (Fig 3E-F, S3E). These results provide a causal relationship between the lack of ORP5 and the perturbation of LD formation and implicate ORP5B in the phenotype.

To address whether the ER-Mito contact sites could play a role in LD formation, we decided to perform similar experiments in cells where these contacts are disrupted. PTPIP51 overexpression increases ER-Mito contact sites, while its knockdown significantly reduces these contacts (De Vos et al., 2012; Stoica et al., 2014) We performed PTPIP51 KD in cells delipidated for 3 days (Fig S4A) and loaded the cells with FA^568^ for 15 min to trigger the formation of LDs. We found a dramatic decrease in the number of FA^568^–positive LDs in the PTPIP51 depleted cells (more than 90%) as compared to control cells (Fig S4B-C).

To test whether ORP5 requires intact ER-Mito contacts to regulate LD biogenesis, we decided to test whether ORP5 overexpression could compensate for the loss of PTPIP51. We performed a rescue experiment in cells depleted for PTPIP51 by overexpressing EGFP-ORP5B and using EGFP-Sec61β as a control. Cells were delipidated for 3 days, loaded with FA^568^ for 1h, and analyzed by confocal microscopy (Fig 3G). Remarkably, ORP5B overexpression, similarly to EGFP-Sec61β, could not rescue the decrease in LDs observed upon PTPIP51 depletion (Fig 3G-H, Fig S4D). These data indicate that the maintenance of the integrity of ER-Mito contact sites is required to ensure ORP5/8-dependent proper LD formation and uncover a novel role of MAMs as key hotspots for LD formation.

### ORP5 is recruited to LDs emerging from MAM subdomains and to pre-existing LDs

To characterize the dynamics of ORP5 localization at MAM-LD contact sites, we performed confocal live-cell imaging by spinning disk microscopy in HeLa cells. Cells transfected with EGFP-ORP5B and Mito-BFP were treated with FA^568^ (at 2 min) and imaged for 12 min. Immediately after the addition of FA^568^, ORP5B was recruited to MAM subdomains where de-novo LDs formed and became visible within 8 minutes following the fatty acid addition (Fig 4A-B, Fig S5A, and movie S1-2).

**Figure 4.**
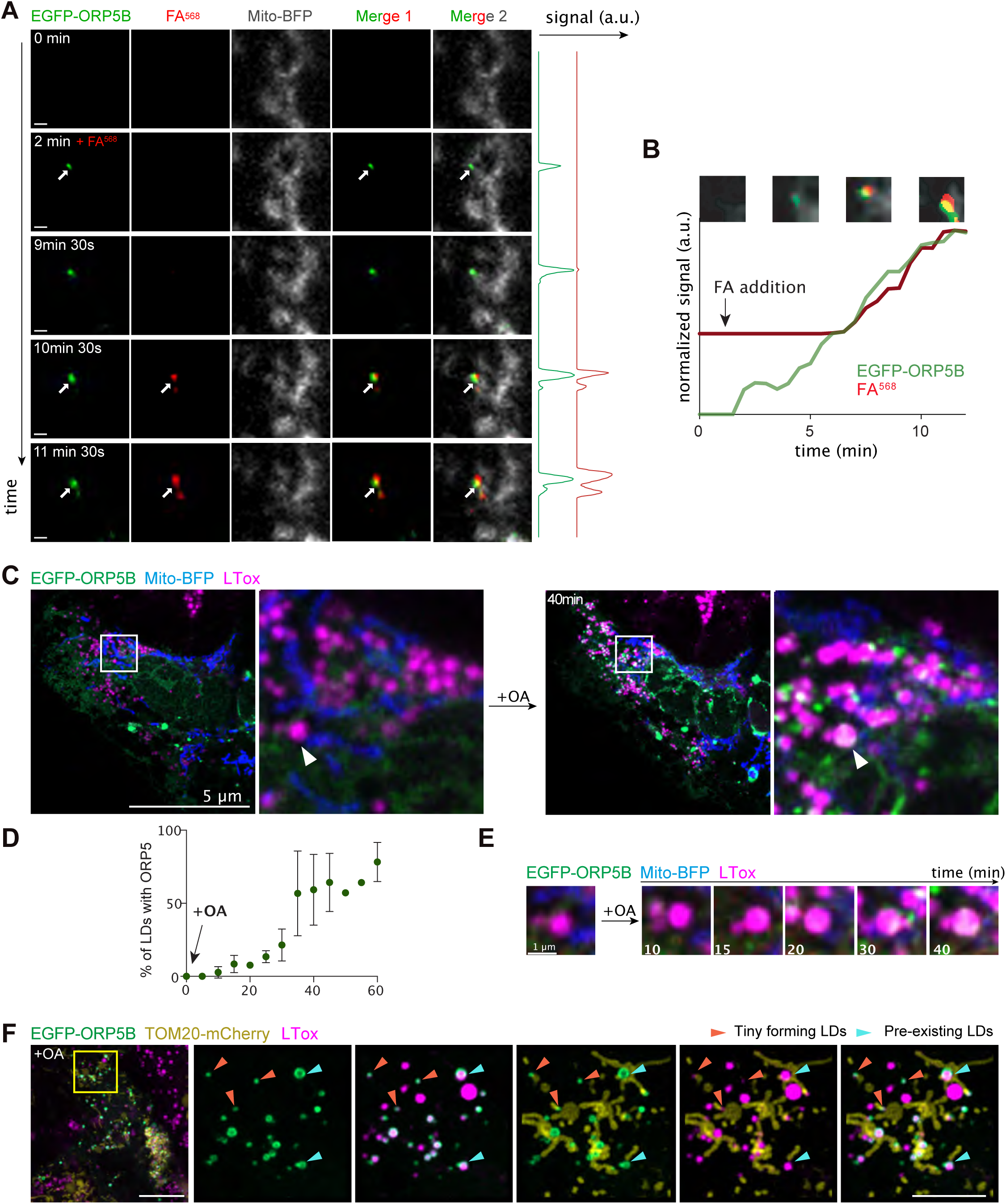
ORP5 specifically localizes to ER subdomains where LDs originate and also to the preexisting lipid droplets. **A)** Zoom of spinning video snapshots of HeLa cells expressing EGFP-ORP5B (green) and Mito-BFP (grey). After 2 min of acquisition, the cells were treated with FA^568^ at 1µM. Arrows indicate ORP5-labeled MAM-LD contacts associated to mitochondria. Full cell view in supplementary figure 5A. Scale bar, 1 µm. **B)** Full time course analysis of the intensity changes for ORP5B (green) and FA^568^ (red) over time. **C)** Example of an Airyscan video snapshots of Huh7 cells expressing EGFP-ORP5B (green), RFP-Sec22b (red, shown in supplementary Figure 5B) and Mito-BFP (blue) before and after 40 min of 200 μM OA treatment. The lipid droplets were stained using LTox Deep Red (purple). Arrowheads indicate absence or presence of ORP5B at MAM-LD contacts before and after OA treatment, respectively. Full sequence in supplementary Figure 5B. Scale bar =10 µm (entire cell) or 5 µm (zoom) **D)** Quantification of the number (%) of LDs with EGFP-ORP5 over the indicated time points. **E)** Time course of ORP5 recruitment to a large pre-existing LDs depicted by the white arrowhead in the Huh7 cell in (C). Scale bar 1 µm. **F)** Representative Airyscan snapshot of Huh7 cells expressing EGFP-ORP5B (green) and TOM20- mCherry (yellow), staining mitochondria, after 1h30min of 200 μM OA treatment. The lipid droplets were stained using LTox Deep Red (purple). Scale bar, 10 µm (entire cell) or 5 µm (zoom). Orange arrowheads indicate tiny emerging LDs, light green arrowheads indicate pre-existing LDs.

To corroborate these data in another cell model relevant for LD physiology, we examined the dynamics of ORP5 recruitment at MAM-LDs contact sites in human hepatocytes Huh7 by Airyscan microscopy in live cells (Airyscan). Huh7 cells were transfected with EGFP-ORP5B, RFP-Sec22b, and Mito-BFP (Figure 4C), or EGFP-ORP5B, TOM20-mCherry, and Mito-BFP (Figure 4F). We then imaged a cell and added OA for 1hr to induce TG synthesis and de novo LD formation.

At 20-25 minutes from OA addition, ORP5 started to be enriched in ER subdomains often corresponding to MAMs, in contact with both LDs and mitochondria (Figure S5B-S5C). The ER-protein Sec22b also localized at the MAM where ORP5 was recruited (Figure S5B), confirming that these structures were indeed ER. Even after 1hr from the induction of LD formation, Sec22b maintained its reticular localization, but it was not enriched at MAM (Fig S5C). In contrast, at 50min-1hr from the OA addition, ORP5 was still strongly at MAM-LDs contact sites (Fig S5B, S5C-D). OA triggered the strong redistribution of ORP5 from the reticular ER to MAM-LD contact regions (Figure 4C-4D, S5B, S5C-D) and the fraction of LDs that are positive for ORP5 increased during feeding (Figure 4D). Tiny LDs, newly emerging, had a strong ORP5 signal (Figure 4E), close to mitochondria also. This observation is consistent with the early recruitment of ORP5 at the MAM sites where LDs assembled in the HeLa cells (Figure 4A, S5A). Thus, both HeLa and Huh7 hepatocyte cells indicate that ORP5 is involved in orchestrating the early stage of LD assembly. However, OA treatment induced targeting of ORP5 not only to mitochondria-associated newly-formed tiny LDs, but also to the preexistent larger ones (Figure 4C-4F, S5B-C), suggesting a role of ORP5 in LD biogenesis but also in their maintenance at MAMs, possibly by regulating lipid fluxes toward them to fit local cellular needs.

### ORP5 is recruited to PA-enriched MAM subdomains

The ORD domain of ORP5 is involved in LD binding, while its PH domain is not required (Fig 1 and 2), as previously shown (Du et al., 2020). The CC domain, which is poorly characterized and sits just before the PH domain, has been proposed to play a role in the association of ORP5 with the plasma membrane (Ghai et al., 2017). We asked whether the ORP5 CC domain could also be relevant for the targeting of MAM-LDs and LD binding. To answer this question, we analyzed by confocal microscopy the localization of the EGFP-tagged ORP5 deletion mutant lacking the CC domain (aa 96-116) (EGFP-ORP5ΔCC) in HeLa cells treated with OA (Fig 5A). As awaited, the deletion of CC in ORP5A led to the loss of ER-PM contacts localization, indicating that the CC might be involved in the binding of PM proteins or lipids. EGFP-ORP5ΔCC localized to the reticular ER in all transfected cells and, only in very few cells (about 10%), it was detected at MAM-LDs contact sites. Also, in these cells, even after OA treatment, ORP5 was not enriched at MAM-LD contacts (Fig 5A). This data strongly supports that the CC domain plays a key role in the association of ORPs with MAM/LDs.

**Figure 5.**
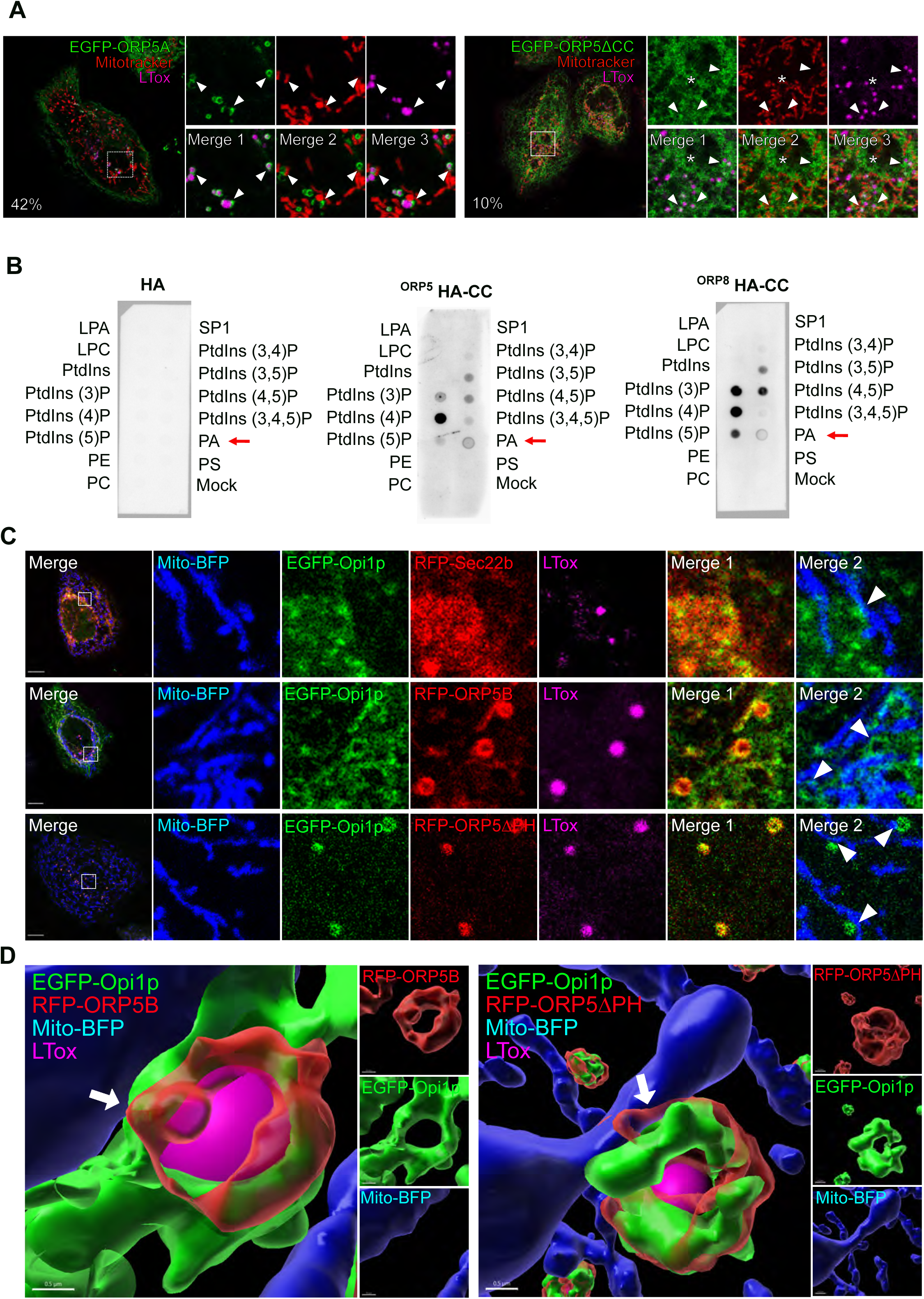
ORP5 localizes to LDs and ER subdomains enriched in phosphatidic acid (PA) **A)** Confocal images (single focal plane of HeLa cells expressing EGFP-tagged ORP5A or ORP5ΔCC (green), treated with OA (300μM) for 2hr. The mitochondria and the LDs were stained with Mitotracker (red) and LTox (purple) respectively. Arrowhead points ORP5-labeled MAM-LD associated to mitochondria and asterisks marks ORP5 localized to reticular ER. Scale bar = 10 µm. **B)** PIP strip overlay assay: PIP strips were incubated with either ORP5-HA CC or ORP8-HA CC or the HA peptide as a negative control and analyzed using the anti-HA antibody. LPA, lysophosphatidic acid; LPC, lysophosphocholine; PtdIns, phosphatidylinositol; PtdIns(3)P; PtdIns(4)P; PtdIns(5)P; PtdIns(3,4)P2; PtdIns(3,5)P2; PtdIns(4,5)P2; PtdIns(3,4,5)P3; PA, phosphatidic acid; PS, phosphatidylserine; PE, phosphatidylethanolamine; PC, phosphatidylcholine; S1P, sphingosine 1-phosphate. **C)** Confocal images (single focal plane) of HeLa cells co-expressing EGFP-Opi1p (green) with Mito-BFP (blue) and either EGFP-Sec22b (red) or EGFP-ORP5B (red) or EGFP-ORP5ΔPH. The LDs were stained with LTox (purple). Arrowheads points enrichment of Opi1p at Mito-MAM-LD contact sites. Scale bar = 10 µm. **D)** 3D reconstruction of cells shown in (D) using IMARIS. Arrows point the MAMs where ORP5B and ORP5ΔPH co-localize with Opi1p at Mito-MAM-LD contact sites. Scale bar = 0.5 µm.

Because the CC domain bears multiple charges, we hypothesized it might bind to charged lipids at MAM/LDs. To assess the lipid-binding specificities of the ORPs CC domains, we performed PIP-strip binding assays. Both ORP5 and ORP8 strongly bound PtdIns(4)P, but ORP8 showed a higher preference for PtdIns(3)P and PtdIns bisphosphates (except for PtdIns(3,4)P, not recognized by the ORPs). Interestingly, ORP5 and ORP8 CCs also bound phosphatidic acid (PA), and, although this binding was quite weak, it was very specific as no other phospholipids were recognized (Fig 5B).

PA is a negatively charged non-bilayer lipid, constantly being made in the ER and essential for the synthesis of phospholipids and TAG, and consequently for LD biogenesis (Gao et al., 2019). We next sought to investigate the presence of PA at MAM-LD contact sites by using a PA-specific probe, the EGFP-tagged Opi1pQ2S-PABD (hereon called EGFP-Opi1p), an improved version of Opi1 for sensing PA. Hela cells were co-transfected with EGFP-Opi1p and either RFP-Sec22b, RFP-ORP5B, or RFP-ORP5ΔPH and analyzed by confocal microscopy. EGFP-Opi1p was detected at RFP-Sec22-positive ER subdomains, and these were often enriched in proximity to mitochondria, revealing the existence of a specific PA pool at MAMs (Fig 5C). When co-expressed with RFP-ORP5B, Opi1p was strongly enriched at the ORP5-marked MAM and MAM-LD contact sites. The enrichment at the latter contact sites was even clearer when Opi1p was co-expressed with RFP-ORP5ΔPH (Fig 5C-D), which had the highest recruitment to MAM-LD contact (Figure 1B, S1A). These data suggest a functional link between PA and ORP5 at MAMs.

### Seipin localizes to MAM-LD contacts in an ORP5-dependent manner

Seipin is an integral ER membrane protein playing a central role in determining where LDs form. Interestingly, seipin binds anionic phospholipids, including PA (Yan et al., 2018). We asked whether seipin could also localize at MAM to facilitate LD formation in proximity to mitochondria. To address this, we co-expressed in HeLa cells a YFP-tagged mouse seipin with Mito-BFP, alone or together with RFP-Sec22b. We then analyzed seipin localization by confocal microscopy (Fig 6A). As expected, seipin colocalized with the ER protein Sec22b. However, a subset of cells where seipin was expressed at lower levels displayed additional enrichments of seipin to ER structures (MAM) in close proximity to mitochondria, which were often also associated with LDs (MAM-LDs). We then asked whether ORP5 could regulate the localization of seipin at MAM. To address this, in parallel experiments, we co-expressed YFP-seipin with RFP-ORP5ΔPH (Fig 6A). Remarkably, we observed an increase in the population of cells showing the enrichment of seipin at MAM-LD contacts (Fig S6A-B).

**Figure 6.**
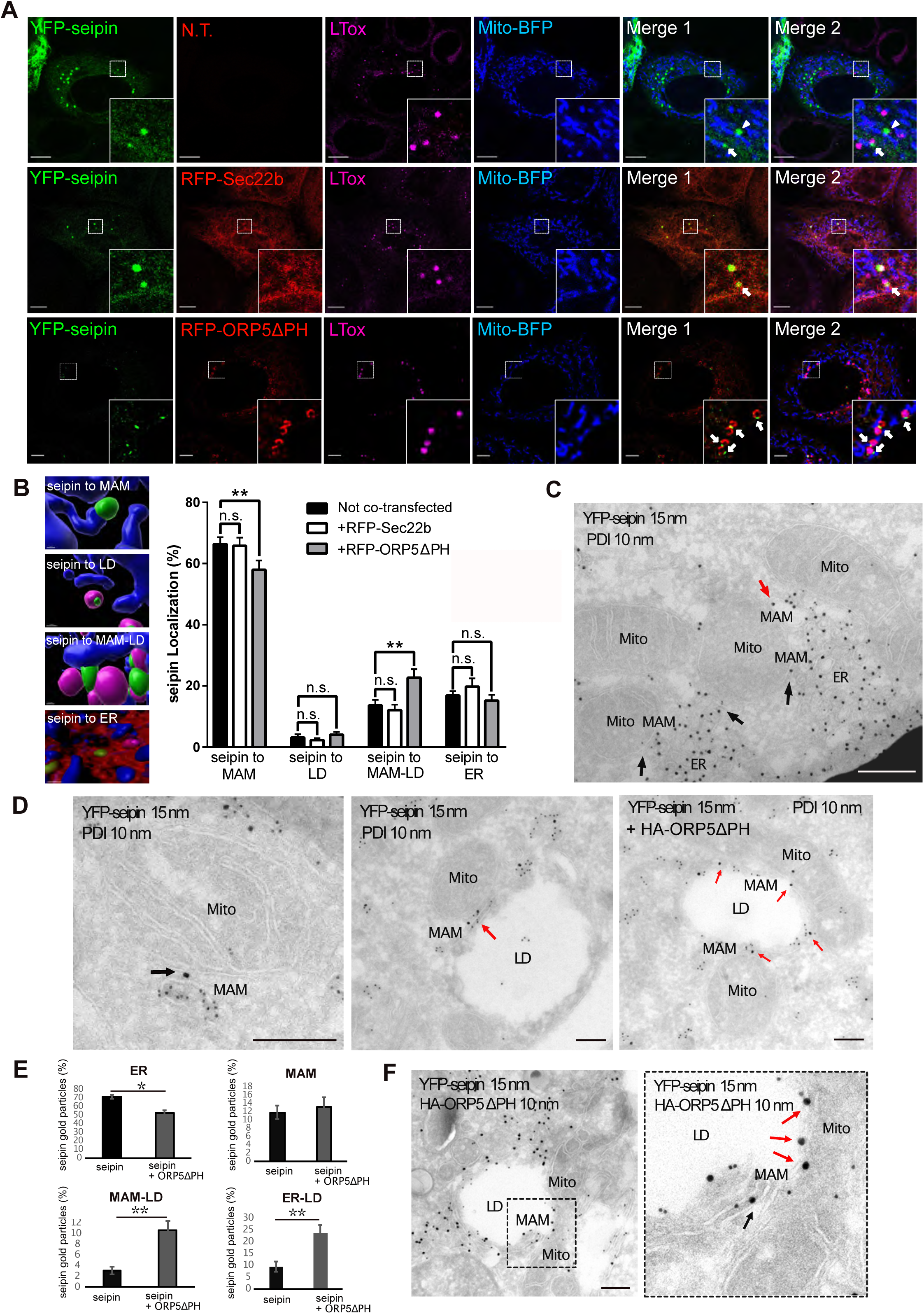
ORP5 over-expression induces an increase of the localization of seipin to MAM-LD contact sites. **A)** Representative confocal images showing a single focal plane of HeLa cells expressing YFP-seipin (green), Mito-BFP (blue) and Sec22b (red) or ORP5ΔPH (red). The LDs were stained LTox (purple). Arrowhead points seipin enrichment at MAM- mitochondria contact sites and arrows mark seipin enrichment at Mito-MAM-LD contact sites. Scale bar = 10 µm. **B)** Analysis of the localization of seipin enrichments in HeLa cells expressing seipin alone or in co-expression with Sec22b or ORP5ΔPH. Representative 3D reconstruction images of the different categories for the classification of the localization. Data are shown as % mean±SEM of cell of n= 56 cells in YFP-seipin (expressed alone), n = 14 cells in YFP-seipin + RFP-Sec22b, and n =40 cells in YFP-seipin + RFP-ORP5ΔPH, (** = p<0.01; Wilcoxon-Mann-Whitney test). **C-D)** Representative images of electron micrographs of ultrathin cryosections of HeLa cells transfected with YFP-seipin and immunogold stained with anti-GFP (15 nm gold) to detect seipin and anti-PDI (10 nm gold) to label the ER lumen. Arrows point at seipin localization at MAM (black arrows) and at MAM-LD contacts (mitochondria-ER-LD) (red arrows). Mito, mitochondria; ER, Endoplasmic Reticulum; MAM, Mitochondria Associated Membranes; LD, Lipid Droplets. Scale bar, 250 nm. **E)** Quantification of the distribution of seipin immunogold particles (15nm). Data is shown as % mean±SEM of cell profiles with n=32 (750 gold particles analysed) in seipin individual expression, and n = 50 (940gold particle) in seipin + ORP5ΔPH co-overexpression. *p<0.001, **p<0.0001, unpaired two-tailed *t*-test. **F)** Electron micrographs of ultrathin cryosections of HeLa cells co-transfected with YFP-seipin and HA-ORP5ΔPH and immunogold stained with anti-GFP (15 nm gold) to detect seipin and anti-HA (10 nm gold) to detect ORP5. The localization of seipin at MAM-LD contacts is increased when co-expressed with ORP5ΔPH (red arrows).

Quantification of the distribution of seipin-positive clusters upon segmentation of the ER, the LD, and the mitochondria by Imaris (Fig 6B) revealed that, when seipin was expressed alone or with Sec22b, most of these clusters corresponded to MAM (about 78%) and only a few (20%) to the reticular ER. A fraction of the seipin-positive MAM (18%) was also closely associated with LDs (corresponding to MAM-LD contacts), and only a minimal amount of seipin-positive ER clusters (2%) was exclusively associated with LDs. When seipin was co-expressed with ORP5ΔPH, we observed a significant increase of Seipin-positive MAM in contact with LD and a significant decrease of seipin-positive MAMs not associated with LD (Fig 6B). However, by confocal imaging, we could not analyze the localization of the entire ER pool of seipin, but only its local enrichment at these “clusters”.

To characterize the distribution of the entire seipin pool at a high-resolution level and to also analyze the ultrastructure of these seipin “clusters”, we performed immuno-EM analysis on ultrathin cryosection (by Tokuyasu method) in HeLa cells expressing YFP- seipin alone or together with HA-ORP5ΔPH, and performed co-immunolabeling of YFP- seipin (15nm gold) and protein disulfide isomerase (PDI) (10nm gold) to stain the ER (Fig 6C-D, Fig S6C). When expressed alone, seipin was mostly observed in the widely distributed reticular ER (70%), but a fraction of seipin (12%) was also found at sites of apposition between MAMs and mitochondria, and a little pool (3%) was detected at MAM-LD contacts closely associated to mitochondria. Few cells displayed accumulation of seipin-positive ER elements in contact with mitochondria that presumably correspond to the “clusters” observed and quantified by confocal. Consistent with our confocal data, these clusters were always found in close connection with mitochondria (corresponding to MAM) and sometimes additionally with LD (corresponding to MAM-LD contacts) (Fig 6C and S6C). Moreover, further corroborating our confocal data, co-expression with ORP5ΔPH increased the pool of seipin at MAM-LD contact sites (3% to 24%) accompanied by an increase of seipin at ER-LD contacts, because of the ORP5-induced expansion of ER on the LD surface (from 10% to 24%). This increase was also accompanied by a significant decrease in seipin abundance at the reticular ER not associated with contact sites (from 70% to 53%) (Fig 6D). Co-localization of seipin and ORP5 at the expanded MAM-LD contact sites was further confirmed by co-immunolabeling of YFP-seipin (15nm gold) and ORP5 (10nm gold) (Fig 6F). Seipin recruitment to ORP5-positive MAM-LD contact sites upon OA was confirmed in Hu7h cells co-expressing seipin–EGFP, RFP-ORP5B, and Mito-BFP (Fig S6D). Overall, these results reveal that seipin localizes at MAM-LD contacts in an ORP5-dependent manner.

### ORP5 role in seipin recruitment to MAM-LD contacts depends on MAM integrity

To further study the role of ORP5, and also of ORP8, in the targeting of seipin to MAM-LD contact sites, we analyzed the localization of seipin in ORP5 and ORP8 KD cells, transfected with YFP-seipin and Mito-BFP and stained by LTox (Fig 7A). Knockdown of ORP5 and ORP8 led to a significant decrease in the cells showing seipin clusters at MAM-LD (Fig S7A) accompanied by an increase in seipin association to the reticular ER (Fig 7A-B). Interestingly, in ORP5-depleted cells, we also detected a general decrease in the seipin pool associated with MAM, while seipin abundance at ER-LD contacts was unchanged in both ORP5- and ORP8-silenced cells (Fig 7A-B). These data reveal a key role of ORP5 and ORP8 in regulating the targeting of seipin to MAM-LD contacts, and a more important role for ORP5 in regulating the general recruitment of seipin to MAMs.

**Figure 7.**
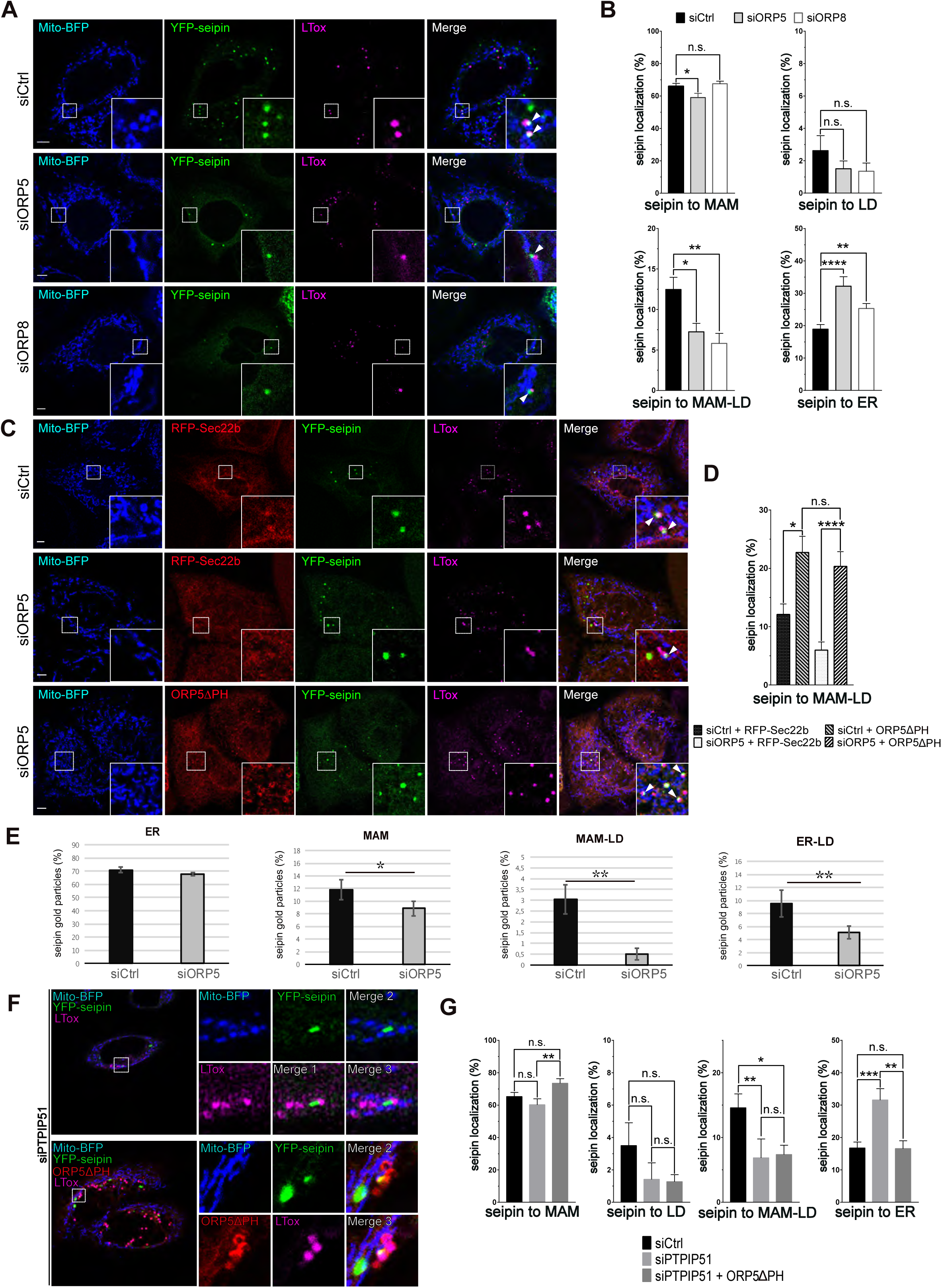
ORP5 affects the localization of seipin in a Mito-MAM contact sites integrity dependent way. **A)** Representative confocal images (single focal plane) of HeLa cells treated with siCtrl or siORP5 or siORP8 RNA oligos. The cells were then transfected with YFP-seipin (green) and Mito-BFP (blue). The LDs were stained LTox (purple). Arrowheads point seipin enrichment at MAM-LD contact sites. **B)** Analysis of the distribution of seipin enrichments in HeLa cells expressing seipin and treated with either siCtrl, siORP5 or siORP8 interfering RNAs. Data are shown as % mean±SEM of cells with n= 64 cells in siCtrl, n = 44 cells in siORP5, and n = 32 cells in siORP8 (* = p<0.05; ** = p<0.01; **** = p <0.0001; Wilcoxon-Mann-Whitney test). **C)** Confocal images (single focal plane) of HeLa cells treated with siCtrl or siORP5 or siORP8 RNA oligos. The cells were then transfected with YFP-seipin (green), Mito-BFP (blue) and either RFP- Sec22b (red) or ORP5ΔPH (red). The LDs were stained LTox (purple). Arrowhead points seipin enrichment at MAM-LD contact sites. **D)** Analysis of the distribution of seipin enrichments to MAM-LD contact sites in HeLa cells treated with siCtrl or siORP5 RNA oligos and then co-transfected with seipin and RFP-Sec22b or siRNA-resistant ORP5ΔPH. Cells were treated with OA (300μM) for 2h before analysis. Data are shown as % mean±SEM of cells. n= 14 cells in siCtrl + Sec22b, n = 17 cells in siORP5 + Sec22b, n = 41 cells in siCtrl + ORP5ΔPH rescue and n = 19 cells in siORP5 + ORP5ΔPH rescue (* = p<0.05; **** = p <0.0001; Wilcoxon-Mann-Whitney test). **E)** Quantification of the distribution of the immunogold particles (15nm) staining seipin in HeLa cells treated with siCtrl or siORP5. Data are shown as % mean±SEM of cell profiles with n=79 (1500 gold particles analysed) in siCtrl, and n = 64 (1800 gold particle) in siORP5. * = p<0.001, ** = p<0.0001, unpaired two-tailed *t*-test. **F)** Representative confocal images (single focal plane) of HeLa cells treated with siPTPIP51 or siCtrl RNAs. The cells were then transfected with YFP-Seipin (green) alone or in co-expression with ORP5ΔPH rescue (red). The LDs were stained with LTox (purple). **G)** Quantification of the distribution of seipin immunogold particles in HeLa cells treated with siCtrl or siPTPIP51 and expressing seipin alone or in co-expression with siRNA-resistant ORP5ΔPH. Data are shown as % mean±SEM of cells with n= 41 cells in siCtrl, n = 25 cells in siPTPIP51, and n =23 cells in siPTPIP51 + ORP5ΔPH (* = p<0.05; ** = p<0.01; *** = p <0.001; Wilcoxon-Mann-Whitney test).

Next, we wanted to rescue the decrease in seipin at MAM-LDs induced by the loss of ORP5. For this purpose, we chose to overexpress ORP5ΔPH in the ORP KD background. We transfected siRNA-resistant ORP5ΔPH, or RFP-Sec22b as a control, in the ORP5 depleted cells and analyzed seipin localization by confocal microscopy (Fig 7C). Re-expression of ORP5ΔPH (detected by immunofluorescence using an antibody against ORP5), but not of RFP-Sec22b, completely rescued the levels of seipin at MAM and even increased the seipin pool at MAM-LD, confirming the specificity of ORP5 activity in controlling seipin localization to MAMs, including those associated with LDs (Fig 7C-D, Fig S7B-C).

To further confirm these observations, we analyzed the entire seipin pool localization at a high-resolution level by performing immuno-EM analysis on ultrathin cryosection in HeLa cells expressing YFP-seipin. We co-immunolabeled YFP-seipin (15nm gold) and the luminal ER protein PDI (10nm gold) (Fig 7E, Fig S7D). A dramatic decrease of seipin localized at MAM-LDs (of about 84%) was observed in ORP5-silenced cells. This decrease was accompanied by a decrease of seipin at ER-LDs and also a slight but statistically significant decrease of seipin at MAMs (Fig 7E). These results further validated the key role of ORP5 in regulating the levels of seipin at MAM and at MAM-LD contacts.

Finally, to assess whether ORP5 requires intact ER-mitochondria contacts to regulate seipin targeting at MAMs, we depleted PTPIP51, which we had found involved in LD biogenesis (see Fig 3G-H and S4). We then analyzed seipin localization by confocal microscopy. PTPIP51 knockdown induced a dramatic decrease (50%) in the number of cells showing seipin localization at MAM-LD contacts (Fig S7E). Also, seipin localization at MAM-LD was greatly reduced (50%), and, in a similar proportion, seipin localization within the reticular ER was increased (Fig 7F-G). Moreover, re-expression of ORP5ΔPH in PTPIP51 depleted cells did not rescue seipin decrease at MAM-LDs, indicating that intact ER-mitochondria contacts are required for ORP5 function in seipin recruitment at MAM-LD contacts.

## DISCUSSION

In this study, we found a novel function of the LTPs ORP5 and ORP8 in regulating LD biogenesis and growth at MAMs. We showed that the ORP5/8 complex localizes at MAM subdomains enriched in PA lipid and where LDs originate. The loss of ORP5/8 impairs LD biogenesis. We then revealed that ORP5 and ORP8 regulate seipin recruitment to the newly-identified MAM-LD contact sites and, importantly, that ER-mitochondria contact sites integrity is required to ensure ORP5/8 function in proper seipin-mediated LD biogenesis.

Amongst all ORP proteins, ORP2 was first identified to localize to LDs to regulate cellular sterol homeostasis (Hynynen et al., 2009; Jansen et al., 2011). Recently, ORP5, but not ORP8, was described at ER-LD contacts (Du et al., 2020). These two proteins have been previously shown to localize at ER-PM and ER-mitochondria contact sites (Chung et al., 2015; Galmes et al., 2016) and exist as a protein complex, mainly at MAMs (Rochin et al., 2021). Yet, ORP5 and ORP8 distribution across all these contact sites remain controversial. Our multiple high-resolution imaging approaches revealed that ORP5 and ORP8 localize and interact at MAM subdomains in contact with LDs (Fig 1-2). Moreover, overexpressed ORP5 induced an expansion of MAMs around LDs due to the strong ability of the ORP5 ORD domain to bind to LDs compared with the ORP8 ORD domain when expressed alone. The ability of ORP8 to enrich in MAM-LD contact sites depends on ORP5 levels: ORP8 enrichment in this site is increased by interaction with ORP5 through its CC domain. These findings highlight the existence of a novel tripartite junctional interface between mitochondria, MAM, and LDs where these two LTPs localize. They also provide novel insights on the role of the CC in ORP5/8, which function was so far poorly understood, in regulating ORP5/8 localization and interaction at MAM-LD contact sites. Interestingly, the knockdown of the ER-mitochondrial tether PTPIP51 leads to a dramatic decrease in newly formed LDs (Fig S4), thus revealing that disruption of ER-mitochondria contact sites perturbs LD biogenesis. This alteration in LD biogenesis was not rescued by the re-expression of ORP5, indicating that ORP5/8 function in LD biogenesis requires intact ER-mitochondria contacts (Fig 3G-H and S4D). Our findings reveal that LD biogenesis and growth occur at ER-mitochondria contact sites, and depend on ORP5/8-activity and the integrity of MAMs.

The metabolic crosstalk between LD and mitochondria is well established for lipid oxidation or storage purposes (Olzmann and Carvalho, 2018; Veliova et al., 2020). Contact sites between LDs and mitochondria form in response to starvation (Herms et al., 2015; Rambold et al., 2015; Wang et al., 2011). During starvation-induced autophagy, DGAT1 (diacylglycerol acyltransferase 1)-dependent LD biogenesis protects mitochondria function by converting fatty acids into TG stored in LDs, to prevent the accumulation of toxic lipids in mitochondria (Nguyen et al., 2017). Recent studies indicate that LD biogenesis can occur at ER subdomains in contact with catabolic organelles, such as the yeast vacuole or the peroxisomes (Hariri et al., 2018; Joshi et al., 2018). Interestingly, DGAT2, one of the two diacylglycerol acyltransferase enzymes converting diacylglycerols into TG, also localizes to MAM (Stone et al., 2009) and might induce the formation of specific LDs originating from MAMs.

Recently, the mitochondrial protein Mitoguardin 2 has been shown to link Mitochondria, LD and ER, to promote de novo lipogenesis in adipocytes from non-lipid precursors (Freyre et al., 2019). Also, interesting, in brown adipocytes, a specific mitochondria subset with restricted dynamics is bound to LDs (Benador et al., 2018). These peridroplet mitochondria support the growth of LDs by providing ATP molecules necessary for TG synthesis (Benador et al., 2018). Also, mitochondria are a site of glycerol biosynthesis, through the Krebs cycle, and could provide glycerol precursors for TG biosynthesis. Based on these knowledge, it may not be surprising that mitochondria transiently or permanently interact with MAMs to support LD biogenesis and maintenance, e.g., by providing ATP or glycerol molecules used to synthesize TG. Currently, molecular mechanisms regulating the functional crosstalk between LD-ER-Mitochondria organelles remain largely unknown.

The ER phospholipid composition is important for LD formation (Ben M’barek et al., 2017; Thiam and Forêt, 2016; Zoni et al., 2021). ORP5/8 may regulate LD biogenesis by regulating the phospholipid composition at MAM. ORP5/8 have been shown to counter-exchange PS and PtdIns(4)P or PtdIns(4,5)P at ER-PM contacts (Chung et al., 2015; Ghai et al., 2017). Recently, ORP5 has been proposed to play a similar role at ER-LD contact sites (Du et al., 2020). However, that ORP5/8 systematically counter exchange PS with PtdIns(4)P is not established. For instance, we recently found that ORP5/8 can transfer PS from the ER to mitochondria at ER-mitochondria contacts despite the lack of PtdIns(4)P on mitochondria. This result suggests that ORP5/8 might carry out multiple lipid transfer activities and that the underlying mechanisms might be different depending on the local lipid composition. In particular, *s*everal pieces of evidence link PA to LD biogenesis (Gao et al., 2019; Pagac et al., 2016) and, interestingly, seipin can bind to PA *in vitro* (Yan et al., 2018). Our confocal and immuno-EM analysis uncovers that seipin localizes at MAM and MAM associated with LDs, in addition to its previously-reported localization at ER-LD contact sites (Salo and Ikonen, 2019; Salo et al., 2019; Szymanski et al., 2007; Wang et al., 2016). This seipin localization to MAM is dependent on ORP5/8, as the knockdown of the latter decreased seipin at MAM-LD junctions. At the same time, the overexpression of an ORP5 variant enriched at MAM-LD induced seipin accumulation at these contact sites (Fig 6-7). This impact of ORP5 function in seipin recruitment to MAM-LD depends on the integrity of the ER-mitochondria contacts. These data identify ORP5 and ORP8 as novel critical players in LD biogenesis by regulating seipin targeting MAMs.

Based on the above observation, one intriguing hypothesis is that ORP5/8 could be involved in PA metabolism at MAMs. ORP5/8 could enrich and cooperate with PA at MAM sites via their lipid transfer activity, e.g., from Mito-to-ER. Indeed, we found that the MAM subdomains where ORP5 localize are enriched in PA lipid (Fig 5), providing the first evidence of the enrichment of PA at MAM. Such acute regulation of local PA levels at MAMs could be necessary for stabilizing the seipin complex, which then better controls LD nucleation and growth by fueling TG from the ER to the newly formed or pre-existing LDs. Such local PA and seipin recruitment by ORP5/8 is here triggered OA induction, which massively redistributes ORP5 to the MAM subdomains in contact with both nascent and pre-existent LDs (Fig 4 and S5). As the CC of ORP5/8 binds to PA *in vitro* (Fig 5), it is possible that in addition to being involved in ORP5-8 interaction, the CC domain of ORP5/8 is also involved in their recruitment to the PA-enriched MAM-LD subdomains, or recruit PA; deletion of the CC decreased ORP5 targeting to MAM-LD contact sites.

Finally, our EM analysis unveils the morphological features of the ER subdomains from which LDs emerge. In cells treated with OA, we observed electron-dense membrane structures, partially invaginated into the LD, that link the LD to the tubular ER elements and that are often found close to mitochondria (Fig 3C). Although their origin was unknown, similar electron-dense structures were recently observed at LD-mitochondria contact sites (Ma et al., 2021). We reveal that these structures correspond to MAM and are strongly decreased in ORP5-depleted cells (Fig 3D). The decrease in the LD population originating from MAM is accompanied by an increase in the LD population still linked to the ER but not in direct contact with mitochondria.

To conclude, our study uncovers an unprecedented role of ORP5 and ORP8 in orchestrating LD biogenesis at MAM. Our findings offer exciting perspectives in a more profound understanding of LD formation in cells and lipodystrophies, and neuronal disorders. Indeed, ORP5/8, seipin, and MAMs, which we now establish to localize in the same subdomains, are critical players of cellular lipid and calcium homeostasis, dysregulated in the onset of the above disorders.

## MATERIAL AND METHODS

### Cell culture and transfection

HeLa cells were cultured in DMEM (Life Technologies) containing GlutaMax (Life Technologies) and supplemented with 10% FBS (Life Technologies), 1% penicillin/streptomycin (Life Technologies), and 1% non-essential amino acids (Life Technologies) at 37°C and 5% CO_2_. For LD biogenesis experiments, HeLa cells were cultured in DMEM (Life Technologies) containing GlutaMax (Life Technologies) and supplemented with 5% lipoprotein-deficient serum FCS (Life Technologies), and 1% non-essential amino acids (Life Technologies) at 37°C and 5% CO_2_ for 72 h, and then treated with BODIPY^TM^ 558/568 (FA^568^) or oleic acid in serum depleted DMEM. For imaging, HeLa cells were seeded in 13-mm glass coverslips. HeLa cells were transfected with the indicated plasmids for 3h in serum depleted medium (Opti-MEM^TM^, ThermoFisher) using lipofectamine 2000 (Life Technologies), according to manufacturer’s protocol. Cells were imaged 16-24hr hours post transfection.

Human hepatocarcinoma cells Huh7, were maintained in High Glucose with stabilized Glutamine and with Sodium Pyruvate Dulbecco’s modified Eagle’s Medium (DMEM) (Dutscher) supplemented with 10% heat-inactivated fetal bovine serum and 1% penicillin/streptomycin (GibcoBRL). The Huh7 cells were transfected with indicated plasmid using Polyethyleneimine HCl MAX (Polysciences) following the manufacturer’s instructions. For the swelling experiments, the Huh7 cells were first transfected with the plasmids and loaded with oleic acid for 24hr. Then, the culture media was next replaced by a hypotonic DMEM culture media, diluted twenty times by water. Cells were then imaged five minutes after the hypotonic medium addition.

### siRNAs oligonucleotides

Transient ORP5, ORP8 and PTPIP51 knockdowns in HeLa cells were performed by transfection of siRNA oligos using oligofectamine (Life Technologies) for 5 hours in serum depleted medium (Opti-MEM^TM^, ThermoFisher), according to manufacturer’s protocol. Cells were imaged 48 hours post transfection.

Double-stranded siRNAs were derived from the following references:

| siRNA | Company, Reference |
| --- | --- |
| OSBPL8 | Dharmacon, J-009508-06<br>(Galmes et al., 2016) |
| OSBPL8 | Dharmacon, J-009508-05<br>(Galmes et al., 2016) |
| OSBPL5 | Dharmacon, J-009274-10<br>(Galmes et al., 2016) |
| OSBPL5 | Dharmacon, J-009274-11<br>(Galmes et al., 2016) |
| PTPIP51 | Dharmacon, J-020973-10-0020<br>(Stoica et al., 2014) |
| Non-targeting | Dharmacon, D-001810-10 |

### Plasmids and cDNA clones

EGFP-ORP5A, EGFP-ORP8, EGFP-ORP5ΔPH, EGFP-ORP5ΔTM, EGFP-ORD5 and EGFP-ORD8 were described in (Galmes et al., 2016). mCh-Plin1 was described in (Ajjaji et al., 2019) and YFP-Seipin was described in (Santinho et al., 2020).GFP-Sec22b and RFP-Sec22b were described in (Gallo et al., 2020). GFP-Sec61β and ssHRP-KDEL were kindly gifted by T. Rapoport (Harward University) and D. Cutler (Schikorski et al., 2007), respectively. Mito-BFP (Addgene plasmid #49151; http://n2t.net/addgene:49151), Seipin-EGFP (Addgene plasmid #129719; http://n2t.net/addgene:129719) and TOM20-mCh (Addgene plasmid #55146; http://n2t.net/addgene:55146) were purchased from Addgene.

#### Generation of the ORP5B variant by mutagenesis

The ORP5B natural variant (Du et al., 2020), partially deleted of its PH domain, was generated by site-directed mutagenesis (Quickchange II-XL, Stratagene). The oligos used were:

EGFP-ORP5BΔ134-201_Fw: CCTTTGGGGCCCTTCAGGCTGTCAGCCA
EGFP-ORP5BΔ134-201_Rv: TGGCTGACAGCCTGAAGGGCCCCAAAGG

#### Generation of the ORP5/8 deletion mutants by mutagenesis

The CC and ORD domains of ORP5 or the TM domain of ORP8 were deleted using site-directed mutagenesis (Quickchange II-XL, Stratagene) to generate EGFP-ORP5ΔCC and EGFP-ORP5ΔORD or EGFP-ORP8ΔTM.

The following primers were used:

EGFP-ORP5ΔORD_Fw: CAGAGGAGAACAAGAGTCTGGAGGACCACAGCCCCTGGGAC
EGFP-ORP5ΔORD_Rv: GTCCCAGGGGCTGTGGTCCTCCAGACTCTTGTTCTCCTCTG
EGFP-ORP5ΔCC 93-123_Fw: CCCACCGCCAGGCCCAGCGTGGTC
EGFP-ORP5ΔCC 93-123_Rv: GACCACGCTGGGCCTGGCGGTGGG
EGFP-ORP8ΔTM_Fw: TATTTTCTGCAACAAAAAGACTAGGGGCCCGGGATC
EGFP-ORP8ΔTM_Rv: GATCCCGGGCCCCTAGTCTTTTTGTTGCAGAAAATA

The EGFP-ORP5ΔCCΔTM and the EGFP-ORP5ΔPHΔTM were generated by deletion of the CC and of the PH in the EGFP-ORP5ΔTM using site-directed mutagenesis (Quickchange II-XL, Stratagene). The oligos used to delete the CC are listed above and the oligos used to delete the PH domain were described in (Galmes et al., 2016).

#### Cloning of the RNAi-resistant ORP5 variants

RNAi-resistant EGFP-ORP5A and EGFP-ORP5B were generated by introducing 4 silent point mutations in the region targeted by the 2 siRNA oligos (#10 and #11) by site-directed mutagenesis (Quickchange II-XL, Stratagene) and the following primers:

RESCUE ORP5_siRNA10_Fw: GGGAAGGTCACCATCGA<u>A</u>TG<u>C</u>GCGAAGAACAACTTCCAGGCC
RESCUE ORP5_siRNA10_Rv: GGCCTGGAAGTTGTTCTTCGC<u>G</u>CA<u>T</u>TCGATGGTGACCTTCCC
RESCUE ORP5_siRNA11_Fw: GAAGCCCAAGGGAATCAAGAA<u>A</u>CC<u>C</u>TACAACCCCATCCTGGGGG
RESCUE ORP5_siRNA11_Rv: CCCCCAGGATGGGGTTGTA<u>G</u>GGTT<u>T</u>CTTGATTCCCTTGGGCTTC

The untagged ORP5ΔPH was generated from the RNAi-resistant EGFP-ORP5ΔPH by excision of the EGFP using the enzymes NheI and HindIII followed by Klenow polymerase treatment and ligation using a T4 DNA ligase.

#### Cloning of HA-ORP5wt and HA-ORP5ΔPH

To generate the HA-ORP5wt, the PCR product, carrying the HA tag at the N-terminus of ORP5, was ligated between AgeI and XhoI in the pEGFP-C1 vector (Clontech) to replace the GFP- with the HA-tag. The oligos used are:

RP5 HA Age1_Fw: 5’GGCGGCACCGGTcgccaccATGTACCCATACGATGTTCCAGATTACGCTatgaaggaggaggccttcctc 3’
RP5 Xho1_Rv: 5’ GGCCTCGAGctatttgaggatgtggttaatg 3’

To generate the HA-ORP5 ΔPH, the PH domain in HA-ORP5wt was deleted by site-directed mutagenesis (Quickchange II-XL, Stratagene) as in (Galmes et al., 2016).

#### Cloning of Opi1 pQ2S-PABd

Opi1 pQ2S-PABd probe was generated by inserting the fragment 113-168 from Opi1p (GeneBank M57383.1) using Yeast DNA (gift from Dr. S. Friant, Université de Strasbourg, France) in *Bgl*II and *Eco*RI digested pEGFP-C1. Specificity for biding PA was establishing using a liposome biding assay as described in (Kassas et al., 2017).

#### Cloning of RFP-ORP5ΔTM and RFP-ORP5B

The insert RFP was recovered from a plasmid mTAG-RFP digested with the enzymes NheI (Fast Digest Thermo)/BsGrI(Fast Digest Thermo) or NheI/XhoI (Fast Digest Thermo). Meanwhile, the plasmids EGFP-ORPΔTM and EGFP-ORP5B were respectively digested with the enzymes NheI/BsrGI and NheI/XhoI to remove the tag EGFP. The insert RFP was then ligated on the plasmid without the tag.

### Antibodies, probes and reagents

Primary antibodies used in this study were following: rabbit anti-ORP5 (SIGMA, HPA038712), mouse anti-ORP8 (Santa Cruz, sc-134409), rabbit anti-PTPIP51 (RDM3, SIGMA, HPA009975), mouse anti-PTPIP51 (FAM82C, SIGMA, SAB1407626), mouse anti-GAPDH (Genetex, GTX627408), rabbit anti-GFP (Invitrogen, A11122) and mouse anti-HA (SIGMA, H3663). Dilutions are detailed in the table below.

Mitotracker red (ThermoFisher) and LipidTox^TM^ (LTox, ThermoFisher) were used as probes for the mitochondrial network and the LDs, respectively, following the manufacturer’s instructions. Additionally, in LD biogenesis experiments, LDs were labeled by BODIPY^TM^ 558/568 (FA^568^, fluorescent tagged oleic acid, Thermofisher) or LD540 (Spandl et al., 2009). Oleic acid-Albumin from bovine serum (OA, SIGMA, 03008) was used to induce LD production.

|  |  |
| --- | --- |
| Rabbit anti-ORP5 | IF 1:300<br>PLA 1:200 |
| Mouse anti-ORP8 | PLA 1:200 |
| Rabbit anti-PTPIP51 | PLA 1:200<br>WB 1:800 |
| Mouse anti-PTPIP51 | PLA 1:200 |
| Mouse Anti-GAPDH | WB 1:10000 |
| Rabbit anti-GFP | IEM 1:100 |
| Mouse anti-HA | IEM 1:500 |
| Mitotracker | 1 μM |
| LipidTOX<sup>TM</sup> (LTox) | 10 μM |
| BODIPY<sup>TM</sup> 558/568 (FA<sup>568</sup>) | 1 μM |
| Oleic Acid (OA) | 300 μM |
| LD450 | 10 μM |

### LD Biogenesis and oleic acid treatment

LD biogenesis was induced in delipidated HeLa cells grown on coverslips by treatment with FA^568^ or OA in serum depleted DMEM. For LD biogenesis time-course experiments and LD biogenesis recue experiments, delipidated HeLa cells (control and knockdown for ORP5, ORP8 or PTPIP51) expressing Mito-BFP alone or Mito-BFP together with EGFP tagged-ORP5 were treated with 1 μM FA^568^ and then fixed in 4%PFA at different times. After fixation, LDs were stained with LTox for 30 min in PBS and mounted for observation. A subset of LD biogenesis time-course experiments were performed in delipidated HeLa cells (control and knockdown for ORP5 or ORP8) treated with 300 μM OA and in which mitochondrial network was labelled with Mitotracker. For all other experiments, non-delipidated HeLa cells were treated with 300 μM OA for 2hr before fixation.

### Confocal Microscopy

Images of immunostained cells or cells expressing fluorescent tagged proteins were acquired on Confocal inverted microscope SP8-X (DMI 6000 Leica). Optical sections were acquired with a Plan Apo 63x oil immersion objective (N.A. 1.4, Leica) using the LAS-X software. Fluorescence was excited using either a 405nm laser diode or a white light laser, and later collected after adjusting the spectral windows with GaAsP PMTs or Hybrid detectors. Images from a mid-focal plane are shown. For Huh7 experiments, images were acquired by the ZEISS LSM800 Airyscan.

### Live-cell Imaging

HeLa cells were seeded on glass bottom ibidi chambers (µ-slide 2 wells) 2 days before imaging. The day after seeding, EGFP-ORP5B and Mito-BFP plasmids were transfected with of lipofectamine 2000/well (Invitrogen), according to the manufacturer’s instructions. Cell imaging was performed on an inverted Nikon Ti Eclipse E microscope coupled with a Spinning Disk (Yokogawa CSU-X1-A1) and cage incubator to control both temperature and CO_2_ (37 °C, 5% CO2). After excitation with a 405nm (Vortran, 100 mW), 491nm (Vortran, 150 mW) and 561 nm laser (Coherent, 100 mW), fluorescence from the different fluorescent compounds was detected with a 40X oil immersion objective (PLAN FLUOR; NA: 1.30; Nikon), an emission filter Quad bandpass 440/40 nm, 521/20 nm, 607/34 nm, 700/45 nm (Semrock) and a Prime 95B sCMOS camera (Photometrics). Images were captured every 30 sec during 15 min. Approximatively 2 min after the start of captures, FA^568^ was added to a final concentration of 1 µM to induce the formation of lipid droplets. For live imaging, Huh7 cells were grown in MatTek 3.5mm coverslip bottom dishes and imaged on the ZEISS LSM800 Airyscan microscope.

### Quantifications

*Colocalization analysis:* for co-localization analysis of fluorescent signals, the acquired images were processed using the JACoP plugin in ImageJ to assess the Pearson’s correlation coefficient. The obtained values, ranging from 0 to 1 (1=max correlation), indicated the association between the signals analysed.

*Organelles/structures count:* LTox- and FA-positive LDs and ORP5 labeled-ER were counted in maximal projection confocal images using ImageJ software.

*Nearest-neighbor distance analysis:* the probability densities of distance distribution in Figure S5D was performed by the Mosaic Interaction analysis plug-in for Fiji (Shivanandan et al., 2013).

*Imaris Analysis: quantification of PLA and seipin spots:* in order to quantify PLA spots and categorize seipin spots in HeLa cells, confocal images were treated with the surface function of the software IMARIS (Bitplane, v9.3.1). For PLA quantification, 3D images (PLA foci identified as “spots”, mitochondria identified as “surfaces) were generated from confocal Z-stack images and the shortest distance between each spot center and the nearest point of the surface or cell object was calculated based on a 3D distance map. Spots objects (PLA dots) with a distance smaller than 380nm from surfaces (mitochondria) objects were considered at a close proximity of these objects. The threshold of 380 nm was used as an estimation of the PLA reaction precision including both primary and secondary antibodies (30nm) plus half the FWHM of the PLA amplification signals (350nm). Similary, 3D segmented images of HeLa cells co-expressing seipin and Mito-BFP, and stained with LTox were generated from z-stack confocal images (seipin, mitochondria and LD identified as “surfaces). Seipin 3D surfaces were then classified into 4 different categories (“seipin to MAM”, “seipin to LD”, “seipin to MAM-LD” and “seipin to ER”) according to their proximity to the labeled compartments using a 3D distance map. The results as showed as the percentage of the category to the total number of the green surfaces.

### 3D Structured Illumination Microscopy (SIM)

Super-resolution light microscopy was performed on a Zeiss ELYRA PS.1 SIM microscope, equipped with a Plan-Apochromat 63x/1.40 NA oil-immersion objective (Carl Zeiss). The illumination patterns of the 488, 561 and 642 nm lasers were projected into the sample. The emitted fluorescence light was detected with an EMCCD camera (iXon 885, Andor Technology). Five phase translations and three rotations of the illumination pattern were recorded at each z-plan and image stacks (120-nm increment along z axis) were acquired. The 3D stacks were then computationally reconstructed with the ZEN imaging software package (algorithm of Heintzmann and Cremer) to generate super-resolution 3D SIM images with twofold extended resolution in the three axes (reconstructed image format = 1904 x 1900 pixels, representing voxels of 0.04 x 0.04 x 0.12 µm).

### Immunofluorescence

Labeling of mitochondrial network with Mitotracker was performed by incubating HeLa cells seeded in glass coverslips with a Mitotracker red 1 μM in Opti-MEM^TM^ (ThermoFisher) for 30 min at 37°C and 5% CO_2_. Cells fixation was carried out by incubation with 4% PFA/PBS for 30 min at room temperature. Coverslips were then washed in PBS and incubated with 50 mM NH4Cl/PBS for 15 min at room temperature. For immunofluorescence, cells were incubated with the primary antibodies diluted in blocking solution (1% BSA / 0.1% Saponin in PBS) for 1 hr at room temperature. Coverslips were thereafter incubated with fluorescently-labeled secondary antibodies (1:500 in BB) for 1hr at room temperature. To label LDs, cells were incubated with 10 μM LTox in 1xPBS for 30 minutes at room temperature. After washing with PBS, coverslips were mounted with Vectashield (Vectro Laboratories) on microscopy slides. Images were acquired on a confocal inverted microscope SP8-X (DMI 6000 Leica). Optical sections were acquired with a Plan Apo 63x oil immersion objective (N.A. 1.4, Leica) using the LAS-X software. Fluorescence was excited using either a 405nm laser diode or a white light laser, and later collected after adjusting the spectral windows with GaAsP PMTs or Hybrid detectors. Images from a mid-focal plane are shown.

### *In situ* proximity ligation assay (PLA)

The protein-protein interactions in fixed HeLa cells were assessed using *in situ* PLA (Duolink^®^SIGMA) according with the manufacturer’s instructions. After fixation, HeLa cells co-expressing Mito-BFP and mCh-Plin1, were incubated with primary antibodies, mouse anti-ORP8 (1:200) plus rabbit anti-ORP5 (1:200), mouse anti-ORP8 (1:200) plus rabbit anti-PTPIP51 (1:200), or rabbit anti-ORP5 (1:200) plus mouse anti-PTPIP51 (1:200), in blocking solution (1% BSA, w/v 0.01% saponin, w/v, in PBS) for 1hr at room temperature. PLUS and MINUS PLA probes (anti-murine and anti-rabbit IgG antibodies conjugated with oligonucleotides, 1:5 in blocking solution) were then incubated with the samples for 1hr at 37°C. Coverslips were thereafter washed in 1x wash buffer A and incubated with ligation solution (5x Duolink^®^ Ligation buffer 1:5, ligase 1:40 in high purity water) for 30 minutes at 37°C. After the ligation step, cell samples were washed in 1x wash buffer A and incubated with the polymerase solution (5x Amplification buffer 1:5, polymerase 1:80 in high purity water) for 1h40min at 37°C. Polymerase solution was washed out from the coverslips with 1x wash buffer B and 0.01x wash buffer B. Vectashield Mounting Medium (Vector Laboratories) was used for mounting.

### Electron Microscopy Analysis

#### Conventional EM

For conventional EM, cells grown on 13 mm glass bottom coverslips (Agar Scientific) were fixed with 2.5% glutaraldehyde and 2% PFA in 0.1 M cacodylate, 0.05% CaCl_2_ buffer for 24 hours. After several washes with 0.1 M cacodylate buffer, the cells were postfixed with 1% OsO_4_, 1.5% potassium ferricyanide in 0.1M Cacodylate for 1 hour. After several washes with 0.1 M cacodylate buffer and H_2_O, the cells were stained with 0.5% uranyl acetate for 24 hours. After several washes with H_2_O, the cells were dehydrated in ethanol and embedded in Epon while on the coverslips. Ultrathin sections were prepared, counterstained with uranyl acetate and observed under a 80kV JEOL 1400 microscope equipped with a Orius High speed (Gatan) camera.

#### HRP Detection

HeLa cells expressing HRP-KDEL were fixed on coverslips with 1.3% glutaraldehyde in 0.1 M cacodylate buffer, washed in 0.1 M ammonium phosphate [pH 7.4] buffer for 1 hour and HRP was visualized with 0.5 mg/ml DAB and 0.005% H_2_O_2_ in 0.1 M Ammonium Phosphate [pH 7.4] buffer. Development of HRP (DAB dark reaction product) took between 5 min to 20 min and was stopped by extensive washes with cold water. Cells were postfixed in 2% OsO_4_+1% K_3_Fe(CN)_6_ in 0.1 M cacodylate buffer at 4°C for 1 hour, washed in cold water and then contrasted in 0.5% uranyl acetate for 2 hours at 4°C, dehydrated in an ethanol series and embedded in epon as for conventional EM. Ultrathin sections were contrasted with 4% uranyl acetate and observed under a FEI Tecnai 12 microscope equipped with a OneView 4k Gatan camera.

#### Serial sectioning and 3D reconstruction

Ultrathin serial sections (50nm) of HRP-KDEL-transfected HeLa cells were cut and put on slot grids (EMS) with formvar (1%) and then contrasted with 4% uranyl acetate. 13 and 14 serial sections for ORP5wt and ORP5ΔPH, respectively, were collected and observed under a 80kV JEOL 1400 microscope equipped with a Orius High speed (Gatan) camera. Image alignment was done with imod and segmentation was done manually by using 3Dmod.

#### Immunogold labelling and quantifications

HeLa cells were fixed with a mixture of 2%PFA and 0.125% glutaraldehyde in 0.1 M phosphate buffer [pH 7.4] for 2 hours, and processed for ultracryomicrotomy as described previously (Slot and Geuze, 2007). Ultrathin cryosections were single- or double-immunogold-labeled with antibodies and protein A coupled to 10 or 15 nm gold (CMC, UMC Utrecht, The Netherlands), as indicated in the legends to the figures. Immunogold-labeled cryosections were observed under a FEI Tecnai 12 microscope equipped with a OneView 4k Gatan camera. For the quantification of the distribution of seipin immunogold labeling on ultrathin cryosections, gold particles (15nm) were counted on acquired micrographs of randomly selected cell profiles (the number of cell profiles and of gold particle is indicated in the figure legends). All data are presented as mean (%) ±SEM of three technical replicates.

### Interfacial tension measurements

Interfacial tension measurements were performed using a drop tensiometer device designed by Teclis Instruments (Tracker, Teclis-IT Concept, France) to measure the interfacial tension of oil – water interfaces. In our experiments, the pendant drop is the triolein lipid phase, formed in the aqueous HKM buffer. The triolein – water interface stabilizes at ∼34± 1 mN/m. Adsorption of ORD5/8 translated into a decrease in tension, as the protein got recruited to the oil-water interface. Throughout the adsorption kinetics to either a triolein – water interface, the drop area was constant.

### PIP-Strip

PIP Strips membranes (Echelon Biosciences) were blocked with 3% BSA FFA dissolved in phosphate-buffered saline (PBS) containing 0.1% Tween 20 (3% BSA FFA PBS-T) at room temperature for 60 min.

Blocked PIP Strips were incubated with the same buffer containing the ^ORP5^HA-CC, ^ORP8^HA-CC or HA peptide (negative control) at final concentration 0.35 µM for 1h. The PIP strips were then washed and bounded proteins were detected with Rabbit anti HA antibody.

### Statistical Analysis

The Kolmogorov-Smirnov test was used for the tensiometer studies. The Wilcoxon-Mann-Whitney test was used for the quantification analysis of seipin by confocal imaging. The unpaired two-tailed *t*-test for all the other experiments.

## AUTHOR CONTRIBUTIONS

FG and ART designed the work. VG, VC and MO performed and analyzed most of the confocal imaging experiments. VG and MO performed the live cell imaging experiments. VC performed and analyzed the LD biogenesis experiments and the PLA. VG performed the seipin localization analysis by Imaris. MO performed the imaging on Huh7 and the PIP strip experiments. CS performed and analyzed some of the LD biogenesis experiments. AH, VG and VC generated ORP5 mutants and performed confocal analysis of their localization. AH and FG performed and analyzed the electron microscopy experiments. CB performed the serial sectioning-3D EM. OF performed the structured illuminated microscopy. NV generated and provided tools for PA imaging analysis. KBM performed the interfacial tension measurements. NEK assisted in western blot analysis, electron microscopy and generated some of the constructs for mammalian cell expression. FG and ART wrote the manuscript and all authors contributed to improve the manuscript.

## CONFLICT OF INTERESTS

The authors declare that they have no competing interests.

## DATA AVAILABILITY SECTION

This study includes no data deposited in external repositories.

## SUPPLEMENTARY FIGURES AND LEGENDS

**Figure S1.**
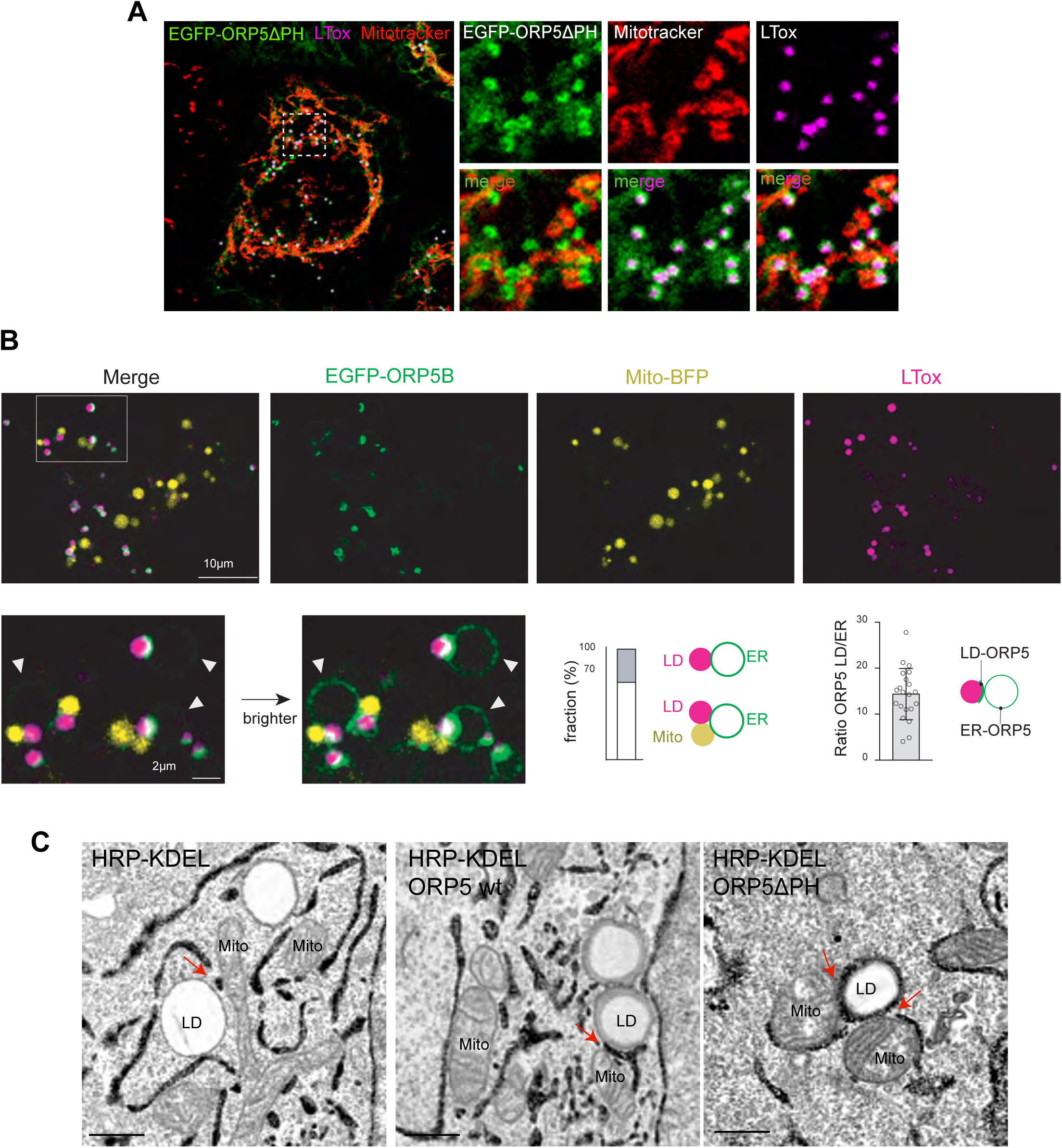
ORP5 localizes to MAM subdomains in contact with LD. **A)** Representative confocal image (single focal plane) of a HeLa cell expressing ORP5ΔPH (green) and treated with Mitotracker (red) and LTox (purple) to label mitochondria and lipid droplets (LD), respectively. Scale bar 10 μm. **B)** Live imaging of EGFP-ORP5 (green), Mito-BFP (yellow) and LDs (purple) ten minutes after swelling Huh7 cells. Zoom area is shown below, normal and enhanced EGFP-ORP5 signal to visualize its ER membrane signal (indicated by arrowheads). The bar graphs, from left to right, show (left) the percentage of ER-LD vs. ER-LD contacts events (n=100) and (right) the signal ratio of EGFP-ORP5 intensity at the ER-LD contacts to the ER membrane (n=20), as depicted by the illustrating image. **C)** Electron micrographs of HeLa cells overexpressing HRP-KDEL alone or together with either EGFP-ORP5A or EGFP-ORP5ΔPH after OA treatment (300 μM for 2hr), and their 3D representation. Scale bar, 250nm

**Figure S2.**
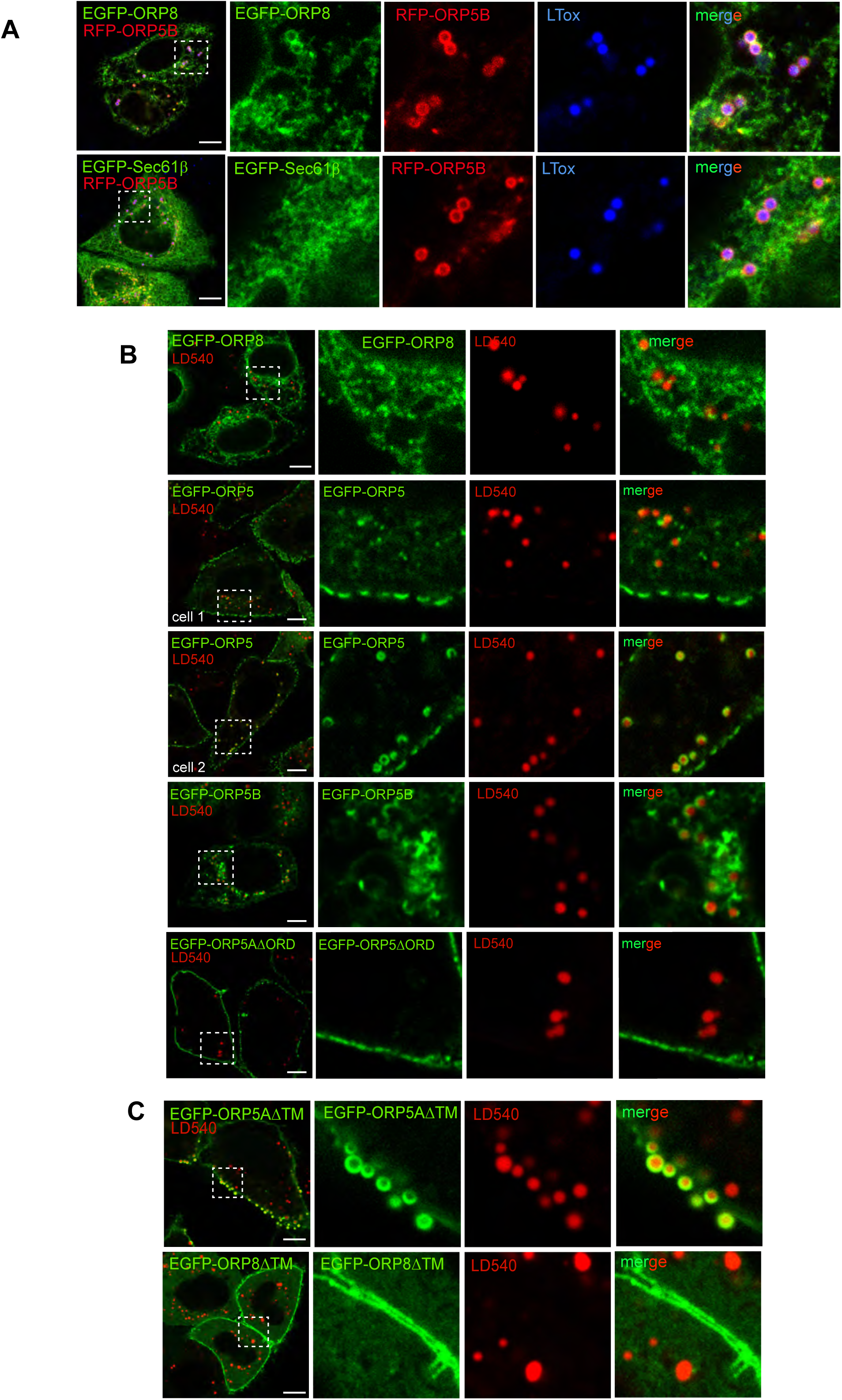
Localization of ORP8 and ORP5 and of their mutants at ER-LD contacts. **A)** Representative confocal images (single focal plane) of HeLa cells co-expressing either EGFP-ORP8 or RFP-Sec61β (green) and RFP-ORP5B (red) and stained with LTox (blue). Scale bar 10 μm. **B)** Confocal images (single focal plane) of HeLa cells expressing EGFP-ORP8, EGFP-ORP5A, EGFP-ORP5B or EGFP-ORP5AΔORD (green) and treated with LD450 (red). Scale bar 10 μm. **C)** Confocal images (single focal plane) of HeLa cells expressing EGFP-ORP5AΔTM or EGFP-ORP8ΔTM (green) and treated with LD450. Scale bar 10 μm.

**Figure S3.**
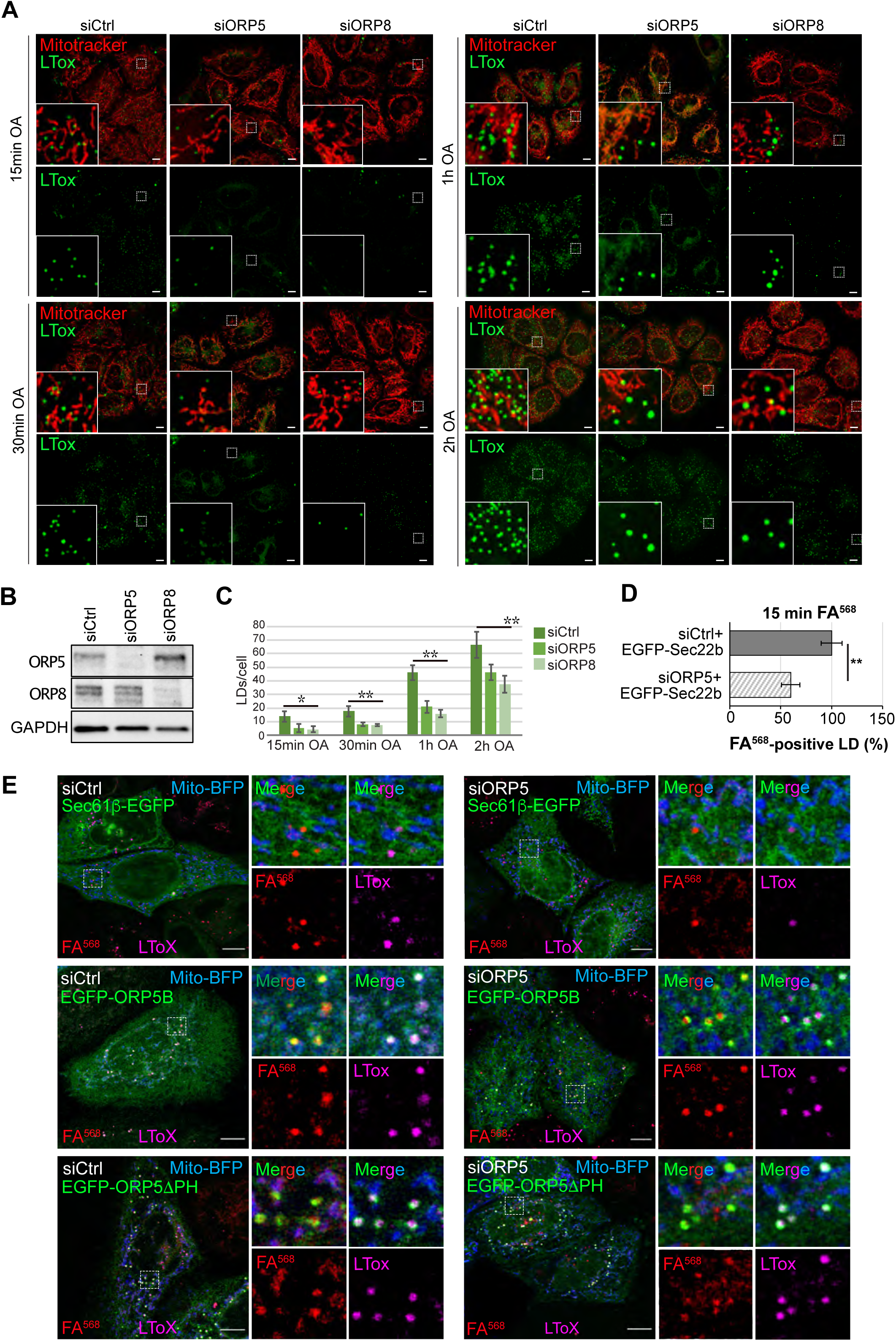
ORP5 and ORP8 depletion impairs LD formation in HeLa cells. **A)** LD biogenesis time-course. Confocal (single focal plane) images of control (siCtrl), and ORP5 (siORP5) or ORP8 (siORP8) siRNA-treated HeLa cells, delipidated for 72h, and incubated with OA (300 μM) for 15 min, 30 min, 1h and 2h. Cells were also stained with Mitotracker (red) and LTox (green). Scale bar, 10 μm. **B)** WB analysis showing ORP5, ORP8 and GAPDH levels in protein lysates from Ctrl, ORP5 and ORP8 knockdown HeLa cells. **C)** Quantification of the number of LTox-positive LDs in siCtrl, siORP5 or siORP8 cells in the indicated times after OA delivery. Data are shown as mean±SEM of n= 30 cells. *p<0.01, **p<0.0001, unpaired two-tailed *t*-test **D)** Analysis of FA^568^-positive LD in siCtrl and siORP5 HeLa cells, priorly delipidated for 72h, and then co-transfected with Mito-BFP and EGFP-Sec22b. Data are show as % of siCtrl treated HeLa cells. n= 27 siCtrl and n=24 siORP5. Bar indicated SEM. **p<0.001, unpaired two-tailed *t*-test. **E)** Representative confocal images of control and ORP5 Knockdown delipidated HeLa cells co-expressing Mito-BFP (blue) and Sec61β-EGFP (green), EGF-ORP5B (green), or ORP5ΔPH (green), and treated with FA^568^ (red, 1 μM for 15 min) and LTox (purple). Images represents a single focal plane confocal series, scale bar 10 μm.

**Figure S4.**
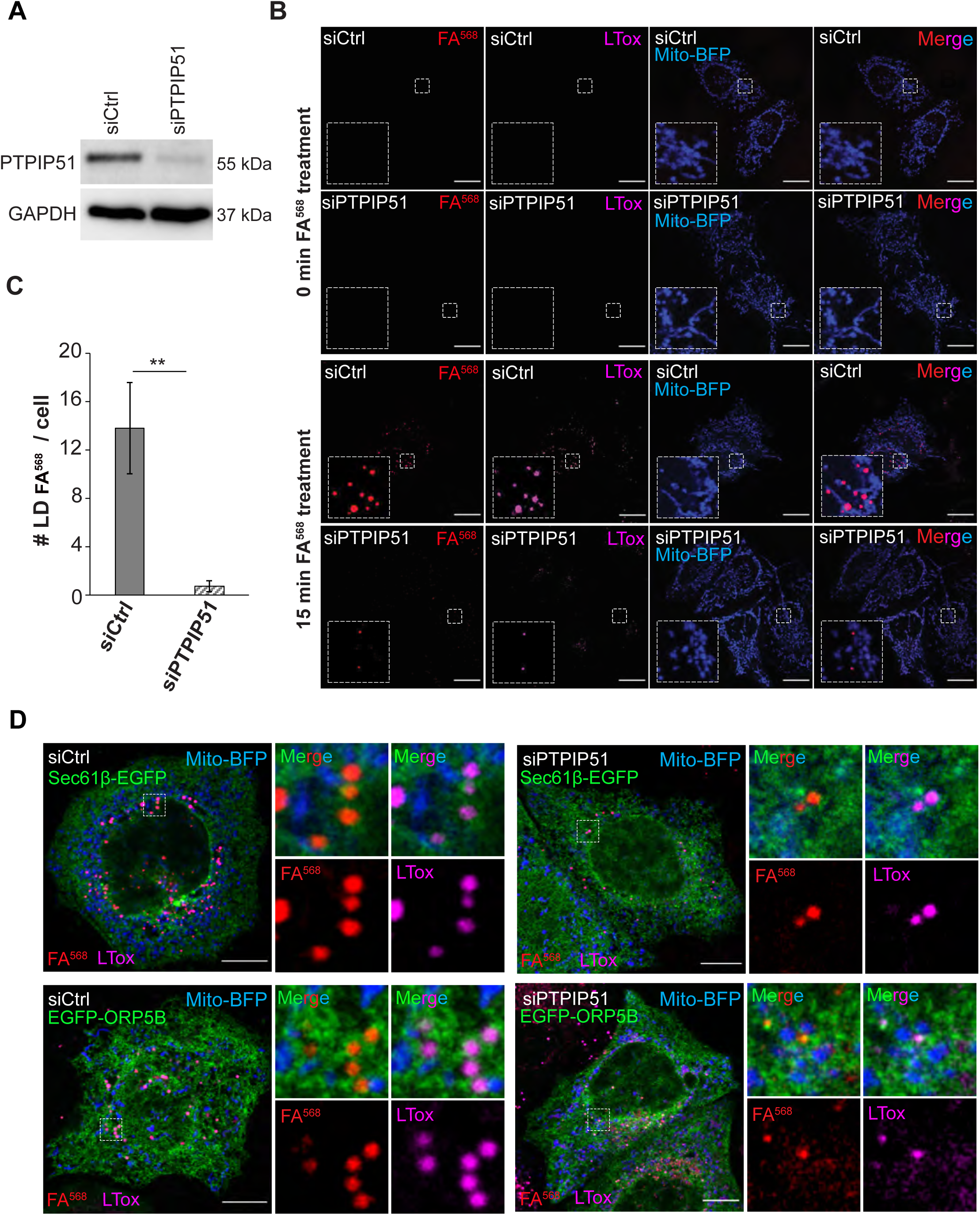
MAM integrity is required for proper LD Biogenesis. **A)** Western blot analysis of the expression of PTPIP51 in siCtrl and siPTPIP51 HeLa cells, showing the efficiency of PTPIP51 knockdown. **B)** Confocal (single focal plane) images of control (siCtrl) and PTPIP51 (siPTPIP51) siRNA-treated HeLa cells, delipidated for 72h, and transfected with Mito-BFP (blue). Cells were treated with FA^568^ (1 μM) for 15 min, and stained with LTox (purple). Scale bar, 10 μm. **C)** Analysis of the number of FA^568^-positive LDs in control and PTPIP51 knockdown cells. Data are show as mean±SEM of n=15 in siCtrl and n=16 in siPTPIP51 cells. ***p<0.001, unpaired two-tailed *t*-test. **D)** Representative confocal images of siCtrl and siPTPIP51 Hela cells, delipidated for 72hr, and transfected with Mito-BFP (blue) and either Sec61β- EGFP (green) or EGF-ORP5B (green). Cells were loaded with FA^568^ (red, 1 μM for 15 min) and stained with LTox (purple) before analysis. Images represents a single focal plane of 3D confocal stacks. Scale bar 10 μm.

**Figure S5.**
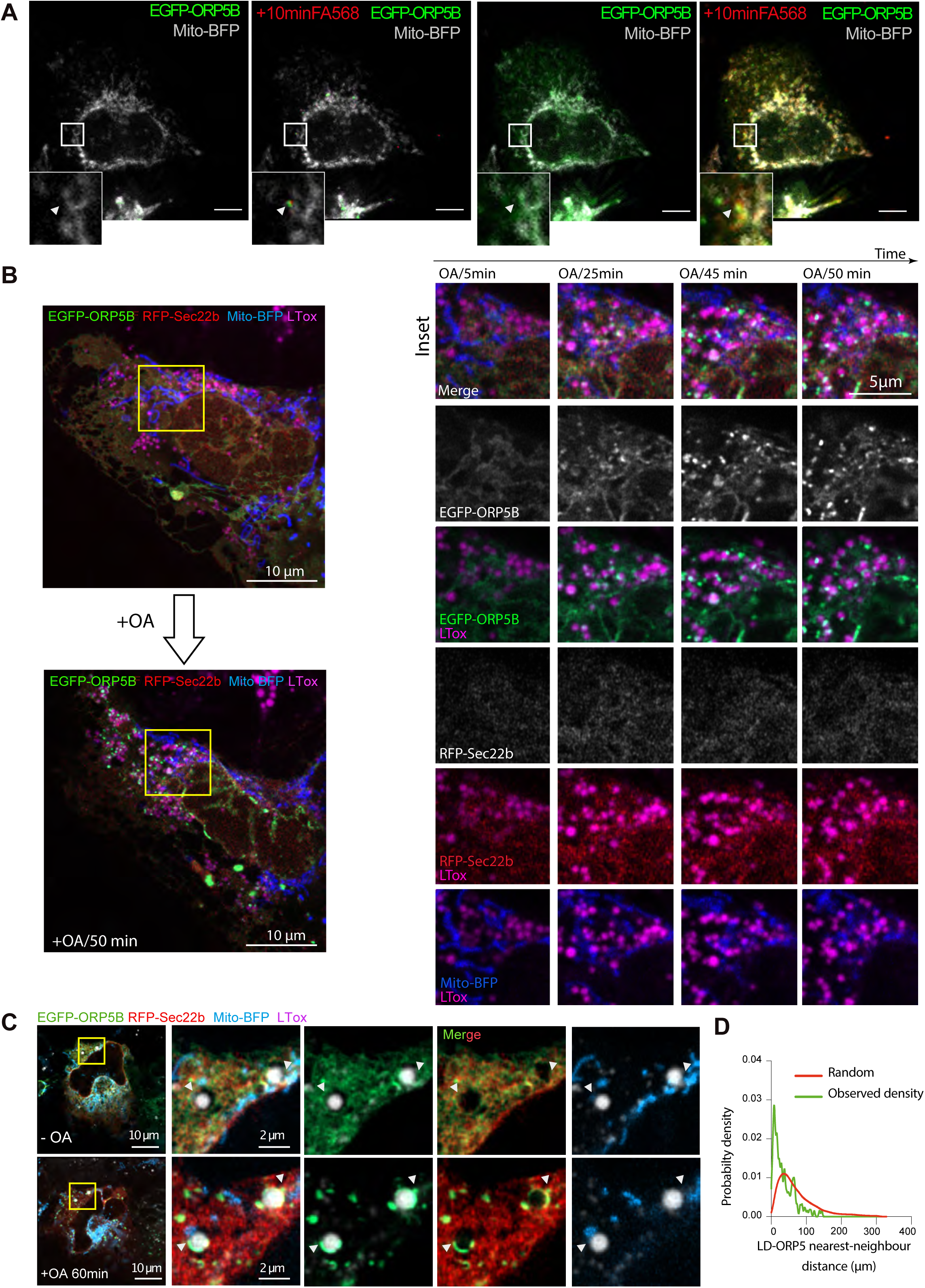
ORP5 redistributes to MAM subdomains where LDs form. **A)** Spinning video snapshots of HeLa cells expressing EGFP-ORP5B (green) and Mito-BFP (grey). After 2min of acquisition, the cells were treated with FA^568^ at 1µM. Left panel shows the cell without brightness and contrast enhancement. Right panel shows the cells with a brightness and contrast treatment to highlight the strongest enrichment of ORP5 and FA^568^ at MAM-LD contacts. Arrowheads indicate EGFP- tagged ORP5 Mito-MAM-LD contacts. Scale bar = 1 µm. **B)** (related to Figure 4C) Airyscan video snapshots of Huh7 cells expressing EGFP-ORP5B (green), RFP-Sec22b (red) and Mito-BFP (blue). Cells were treated with OA to induce the formation of LDs, stained by LTox (purple). Images were taken every 5 min for 50 min. **C)** Another example depicting the recruitment of ORP5 to pre-existing LDs, with mitochondria. Arrowheads indicate EGFP-tagged ORP5 at Mito-MAM-LD contacts. **D)** Line graph shows the distribution of the nearest-neighbor EGFP-ORP5B puncta to LDs in the images of Huh7 cells treated with OA in (B). The observed probability density largely deviates from randomly distributed LD and puncta EGFP-ORP5B.

**Figure S6.**
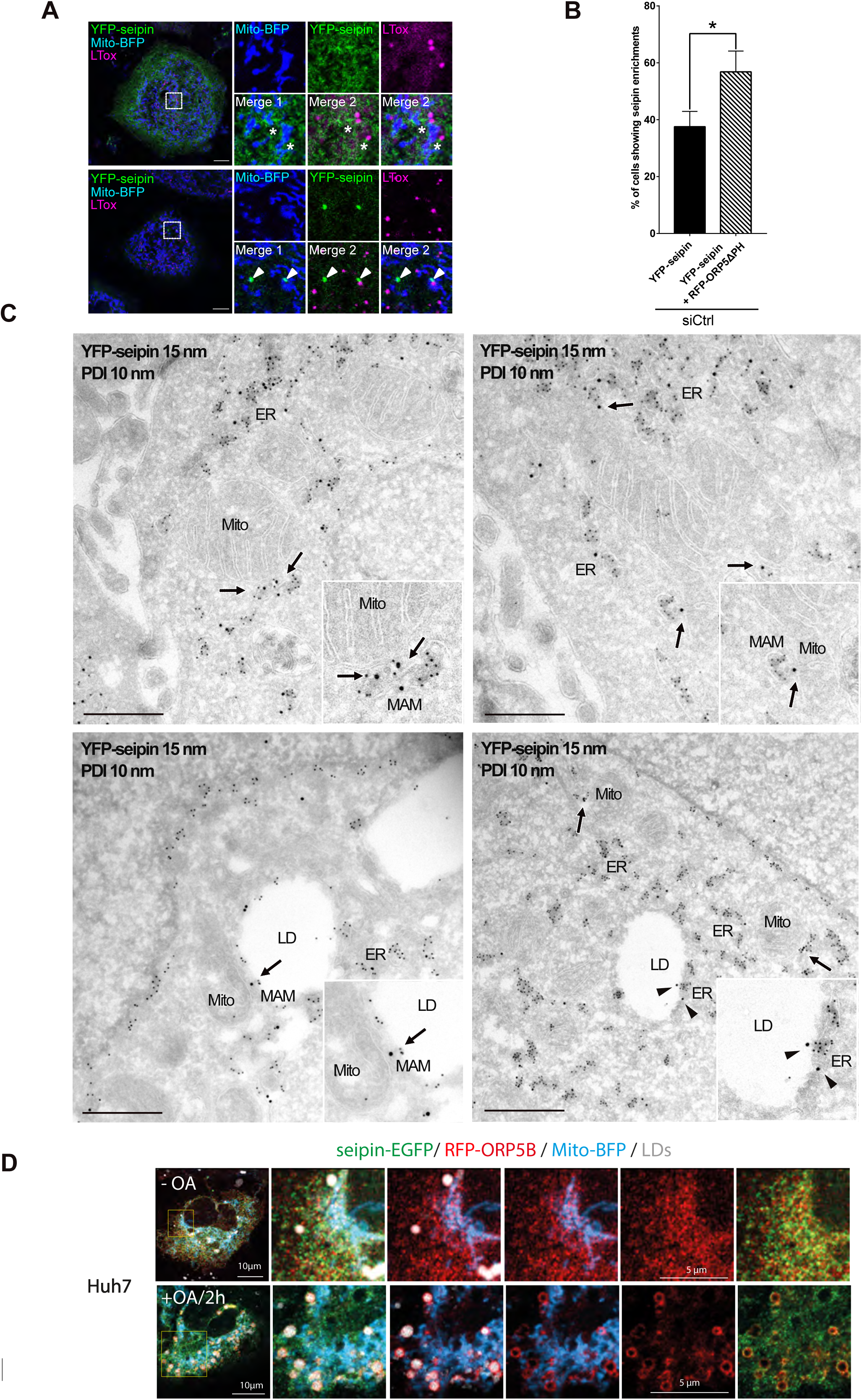
ORP5 over-expression induces an increase of the localization of seipin to MAM-LD contact sites. **A)** Confocal images (single focal plane) of HeLa cells expressing YFP-seipin (green) and Mito-BFP (blue). The LDs were stained LTox (purple). Upper panels show a cell with a reticular staining of seipin. Lower panels show a cell with enrichment of YFP- seipin in small “clusters”. Arrowheads show the enrichment of seipin in “clusters” closely opposed to MAM-LD contact sites. Asterisks show presence of seipin at MAM-LD contacts also in cells where it has a reticular distribution. **B)** Quantification of the % of cells showing seipin enrichment at MAM-LD in cells expressing seipin alone or together with ORP5ΔPH. Data are shown as % mean±SEM of cells. n= 118 cells in siCtrl, n = 121 cells in siCtrl + RFP-ORP5ΔPH. **C)** Representative images of electron micrographs of ultrathin cryosections of HeLa cells transfected with YFP-seipin and immunogold stained with anti-GFP (15 nm gold) to detect seipin and anti-PDI (10 nm gold) to label the ER lumen. Arrows point the localization of seipin at MAM or MAM-LDs. Arrowheads point the localization of seipin at ER-LD contacts. Mito, mitochondria; ER, Endoplasmic Reticulum; MAM, Mitochondria Associated Membranes; LD, Lipid Droplets. Scale bar = 250 nm. **D)** Confocal images (single focal plane) of Huh7 cells expressing seipin-EGFP (green), RFP-ORP5B (red) and Mito-BFP (blue) before and after 2h of OA treatment. The LDs were stained with LTox (white). Scale bar = 10 µm.

**Figure S7.**
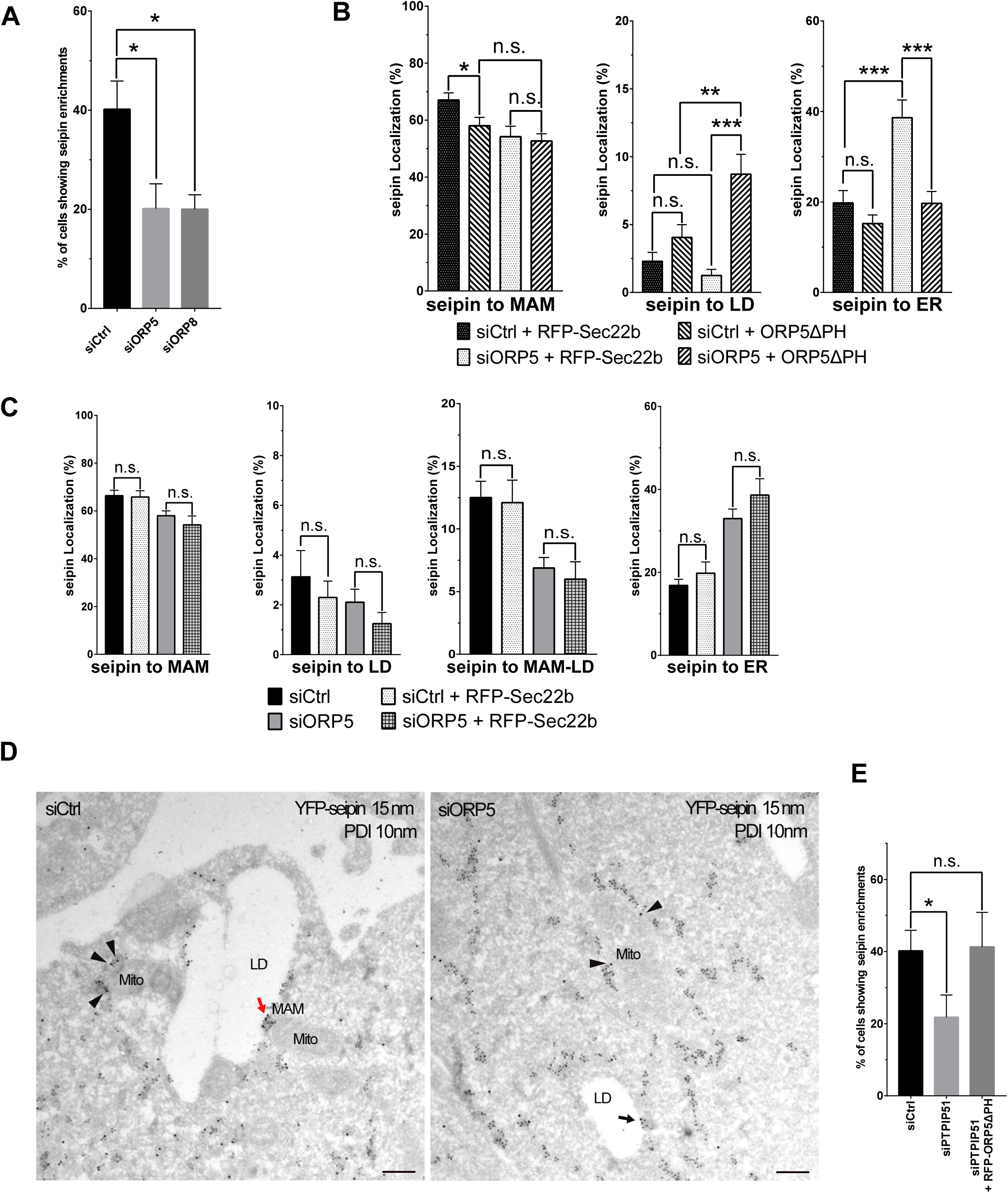
ORP5 affects the localization of seipin in a Mito-MAM contact sites integrity dependent way. **A)** Quantification analysis of confocal data showing the percentage of YFP-seipin– expressing siCtrl or siORP5 or siORP8 cells displaying seipin enrichment at MAM-LD. Data are shown as % mean±SEM of cell of n=118 cells in siCtrl, n = 80 cells in siORP5 and n = 134 cells in siORP8. **B)** Analysis of seipin localization to the indicated compartments (MAM=Mito-MAM contacts, ER-LD=ER-LD contacts, ER=reticular ER) in siCtrl or siORP5 HeLa cells transfected with YFP-seipin and either RFP-Sec22b or siRNA-resistant ORP5ΔPH. Data are shown as % mean±SEM of cells. n= 14 cells in siCtrl + Sec22b, n = 17 cells in siORP5 + Sec22b, n = 41 cells in siCtrl + ORP5ΔPH rescue and n = 19 cells in siORP5+ ORP5ΔPH rescue (* = p<0.05; ** = p<0.01; *** = p <0.001; Wilcoxon-Mann-Whitney test). **C)** Quantitative analysis of the distribution of seipin enrichments in the indicated compartments (MAM=Mito-MAM contacts, ER-LD=ER-LD contacts, MAM-LD=Mito-MAM-LD contacts, ER=reticular ER) in HeLa cells treated with siCtrl or siORP5 and expressing YFP-seipin alone or together with RFP-Sec22b. Data are shown as % mean±SEM of cell of n = 56 cells in siCtrl, n= 14 cells in siCtrl + Sec22b, n = 58 cells in siORP5 and n = 17 cells in siORP5 + Sec22b (n.s. = not significant; Wilcoxon-Mann-Whitney test). **D)** Representative electron micrographs of ultrathin cryosections of HeLa cells transfected with YFP-seipin and immunogold stained with anti-GFP (15 nm gold) to detect seipin and anti-PDI (10 nm gold) to label the ER lumen. Arrows point to seipin localization at MAM-LD (red arrow) or ER-LD (black arrow). Arrowheads point to seipin localization at MAM. Mito, mitochondria; ER, Endoplasmic Reticulum; MAM, Mitochondria Associated Membranes; LD, Lipid Droplets. Scale bar = 250 nm **E)** Quantification of the % of siCtrl or siPTPIP51 cells transfected with YFP-seipin or of siPTPIP51 cells co-transfected with YFP-seipin and RFP-ORP5ΔPH, showing seipin enrichment at MAM-LD. Data are shown as % mean±SEM of cell of n= 118 cells in siCtrl, n = 91 cells in siPTPIP51 and n =58 cells in siPTPIP51 + RFP-ORP5ΔPH.

## SUPPLEMENTARY MOVIES

**Movies S1 and S2: ORP5 localizes to ER subdomains where LDs originate (related to Figure 4)**

Time-lapse of live HeLa cell co-expressing fluorescent proteins Mito-BFP (grey) and EGFP-ORP5B (green). After 2 min of acquisition, the cells were treated with FA^568^ at 1 µM. Movie S1 shows the entire cell and Movie S2 shows a zoom of a region of the HeLa cell.

## Notes

### Competing Interest Statement

The authors have declared no competing interest.

## REFERENCES

Adeyo, O., Horn, P.J., Lee, S., Binns, D.D., Chandrahas, A., Chapman, K.D., and Goodman, J.M. (2011). The yeast lipin orthologue Pah1p is important for biogenesis of lipid droplets. Journal of Cell Biology 192, 1043–1055.

Ajjaji, D., Ben M’barek, K., Mimmack, M.L., England, C., Herscovitz, H., Dong, L., Kay, R.G., Patel, S., Saudek, V., Small, D.M., et al. (2019). Dual binding motifs underpin the hierarchical association of perilipins1-3 with lipid droplets. Molecular biology of the cell 30, 703–716.

Ben M’barek, K., Ajjaji, D., Chorlay, A., Vanni, S., Forêt, L., and Thiam, A.R. (2017). ER Membrane Phospholipids and Surface Tension Control Cellular Lipid Droplet Formation. Dev. Cell 41, 591–604.e7.

Bi, J., Wang, W., Liu, Z., Huang, X., Jiang, Q., Liu, G., Wang, Y., and Huang, X. (2014). Seipin promotes adipose tissue fat storage through the ER Ca2+-ATPase SERCA. Cell Metabolism 19, 861–871.

Choudhary, V., El Atab, O., Mizzon, G., Prinz, W.A., and Schneiter, R. (2020). Seipin and Nem1 establish discrete ER subdomains to initiate yeast lipid droplet biogenesis. Journal of Cell Biology 219.

Chung, J., Torta, F., Masai, K., Lucast, L., Czapla, H., Tanner, L.B., Narayanaswamy, P., Wenk, M.R., Nakatsu, F., and De Camilli, P. (2015). PI4P/phosphatidylserine countertransport at ORP5-and ORP8-mediated ER–plasma membrane contacts. Science 349, 428–432.

Chung, J., Wu, X., Lambert, T.J., Lai, Z.W., Walther, T.C., and Farese Jr, R.V. (2019). LDAF1 and Seipin Form a Lipid Droplet Assembly Complex. Developmental Cell.

De Vos, K.J., Morotz, G.M., Stoica, R., Tudor, E.L., Lau, K.F., Ackerley, S., Warley, A., Shaw, C.E., and Miller, C.C. (2012). VAPB interacts with the mitochondrial protein PTPIP51 to regulate calcium homeostasis. Human molecular genetics 21, 1299–1311.

Du, X., Zhou, L., Aw, Y.C., Mak, H.Y., Xu, Y., Rae, J., Wang, W., Zadoorian, A., Hancock, S.E., Osborne, B., et al. (2020). ORP5 localizes to ER-lipid droplet contacts and regulates the level of PI(4)P on lipid droplets. The Journal of cell biology 219.

Fei, W., Shui, G., Gaeta, B., Du, X., Kuerschner, L., Li, P., Brown, A.J., Wenk, M.R., Parton, R.G., and Yang, H. (2008). Fld1p, a functional homologue of human seipin, regulates the size of lipid droplets in yeast. J Cell Biol 180, 473–482.

Fei, W., Shui, G., Zhang, Y., Krahmer, N., Ferguson, C., Kapterian, T.S., Lin, R.C., Dawes, I.W., Brown, A.J., and Li, P. (2011). A role for phosphatidic acid in the formation of “supersized” lipid droplets. PLoS Genetics 7, e1002201.

Galmes, R., Houcine, A., van Vliet, A.R., Agostinis, P., Jackson, C.L., and Giordano, F. (2016). ORP5/ORP8 localize to endoplasmic reticulum–mitochondria contacts and are involved in mitochondrial function. EMBO Reports 17, 800–810.

Gallo, A., Danglot, L., Giordano, F., Hewlett, B., Binz, T., Vannier, C., and Galli, T. (2020). Role of the Sec22b-E-Syt complex in neurite growth and ramification. Journal of cell science 133.

Gao, M., Huang, X., Song, B.-L., and Yang, H. (2019). The biogenesis of lipid droplets: Lipids take center stage. Progress in Lipid Research 100989.

Ghai, R., Du, X., Wang, H., Dong, J., Ferguson, C., Brown, A.J., Parton, R.G., Wu, J.-W., and Yang, H. (2017). ORP5 and ORP8 bind phosphatidylinositol-4, 5-biphosphate (PtdIns (4, 5) P 2) and regulate its level at the plasma membrane. Nature Communications 8, 1–14.

Gluchowski, N.L., Becuwe, M., Walther, T.C., and Farese Jr, R.V. (2017). Lipid droplets and liver disease: from basic biology to clinical implications. Nature Reviews Gastroenterology & Hepatology 14, 343.

Grippa, A., Buxó, L., Mora, G., Funaya, C., Idrissi, F.-Z., Mancuso, F., Gomez, R., Muntanyà, J., Sabidó, E., and Carvalho, P. (2015). The seipin complex Fld1/Ldb16 stabilizes ER–lipid droplet contact sites. J Cell Biol 211, 829–844.

Han, S., Binns, D.D., Chang, Y.-F., and Goodman, J.M. (2015). Dissecting seipin function: the localized accumulation of phosphatidic acid at ER/LD junctions in the absence of seipin is suppressed by Sei1p ΔNterm only in combination with Ldb16p. BMC Cell Biology 16, 29.

Hariri, H., Rogers, S., Ugrankar, R., Liu, Y.L., Feathers, J.R., and Henne, W.M. (2018). Lipid droplet biogenesis is spatially coordinated at ER–vacuole contacts under nutritional stress. EMBO Reports 19, 57–72.

Herker, E., Vieyres, G., Beller, M., Krahmer, N., and Bohnert, M. (2021). Lipid droplet contact sites in health and disease. Trends in Cell Biology.

Joshi, A.S., Nebenfuehr, B., Choudhary, V., Satpute-Krishnan, P., Levine, T.P., Golden, A., and Prinz, W.A. (2018). Lipid droplet and peroxisome biogenesis occur at the same ER subdomains. Nature Communications 9, 2940.

Joshi, A.S., Ragusa, J.V., Prinz, W.A., and Cohen, S. (2021). Multiple C2 domain–containing transmembrane proteins promote lipid droplet biogenesis and growth at specialized endoplasmic reticulum subdomains. Molecular Biology of the Cell 32, 1147–1157.

Kassas, N., Tanguy, E., Thahouly, T., Fouillen, L., Heintz, D., Chasserot-Golaz, S., Bader, M.F., Grant, N.J., and Vitale, N. (2017). Comparative Characterization of Phosphatidic Acid Sensors and Their Localization during Frustrated Phagocytosis. The Journal of biological chemistry 292, 4266–4279.

Khaldoun, S.A., Emond-Boisjoly, M.A., Chateau, D., Carriere, V., Lacasa, M., Rousset, M., Demignot, S., and Morel, E. (2014). Autophagosomes contribute to intracellular lipid distribution in enterocytes. Molecular biology of the cell 25, 118–132.

Klug, Y.A., Deme, J.C., Corey, R.A., Renne, M.F., Stansfeld, P.J., Lea, S.M., and Carvalho, P. (2021). Mechanism of lipid droplet formation by the yeast Sei1/Ldb16 Seipin complex. Nature Communications 12, 1–13.

Magré, J., Delépine, M., Khallouf, E., Gedde-Dahl Jr, T., Van Maldergem, L., Sobel, E., Papp, J., Meier, M., Mégarbané, A., and Lathrop, M. (2001). Identification of the gene altered in Berardinelli–Seip congenital lipodystrophy on chromosome 11q13. Nature Genetics 28, 365.

Pagac, M., Cooper, D.E., Qi, Y., Lukmantara, I.E., Mak, H.Y., Wu, Z., Tian, Y., Liu, Z., Lei, M., and Du, X. (2016). SEIPIN regulates lipid droplet expansion and adipocyte development by modulating the activity of glycerol-3-phosphate acyltransferase. Cell Reports 17, 1546– 1559.

Rao, M.J., and Goodman, J.M. (2021). Seipin: harvesting fat and keeping adipocytes healthy. Trends in Cell Biology.

Rochin, L., Sauvanet, C., Jääskeläinen, E., Houcine, A., Kivelä, A., Ma, X., Marien, E., Dehairs, J., Neveu, J., Le Bars, R., Santonico E., Swinnen J. V., Bernard D., Tareste D., Olkkonen V. M. and Giordano F. (2019). ORP5 regulates transport of lipids and calcium to mitochondria at endoplasmic reticulum-mitochondria membrane contact sites. BioRxiv 695577.

Rochin, L., Monteiro-Cardoso Vera F., Arora A., Houcine, A., Jääskeläinen, E., Kivelä, A., Sauvanet, C., Le Bars R., Marien, E., Dehairs, J., Neveu, J., El Khallouki N., Santonico E., Swinnen J. V., Tareste D., Olkkonen V. M. and Giordano F. (2021). ORP5/8 and MIB/MICOS link ER-mitochondria and intramitochondrial contacts for non-vesicular transport of phosphatidylserine. BioRxiv 695577.

Salo, V.T., and Ikonen, E. (2019). Moving out but keeping in touch: contacts between endoplasmic reticulum and lipid droplets. Current Opinion in Cell Biology 57, 64–70.

Salo, V.T., Li, S., Vihinen, H., Hölttä-Vuori, M., Szkalisity, A., Horvath, P., Belevich, I., Peränen, J., Thiele, C., and Somerharju, P. (2019). Seipin facilitates triglyceride flow to lipid droplet and counteracts droplet ripening via endoplasmic reticulum contact. Developmental Cell 50, 478–493.

Santinho, A., Salo, V.T., Chorlay, A., Li, S., Zhou, X., Omrane, M., Ikonen, E., and Thiam, A.R. (2020). Membrane Curvature Catalyzes Lipid Droplet Assembly. Current Biology 30, 2481–2494.e6.

Schikorski, T., Young, S.M., Jr., and Hu, Y. (2007). Horseradish peroxidase cDNA as a marker for electron microscopy in neurons. Journal of neuroscience methods 165, 210–215.

Schuldiner, M., and Bohnert, M. (2017). A different kind of love - lipid droplet contact sites. Biochimica et biophysica acta 1862, 1188–1196.

Shivanandan, A., Radenovic, A., and Sbalzarini, I.F. (2013). MosaicIA: an ImageJ/Fiji plugin for spatial pattern and interaction analysis. BMC bioinformatics 14, 349.

Slot, J.W., and Geuze, H.J. (2007). Cryosectioning and immunolabeling. Nature protocols 2, 2480–2491.

So\ltysik, K., Ohsaki, Y., Tatematsu, T., Cheng, J., Maeda, A., Morita, S., and Fujimoto, T. (2021). Nuclear lipid droplets form in the inner nuclear membrane in a seipin-independent manner. Journal of Cell Biology 220.

Spandl, J., White, D.J., Peychl, J., and Thiele, C. (2009). Live cell multicolor imaging of lipid droplets with a new dye, LD540. Traffic 10, 1579–1584.

Stoica, R., De Vos, K.J., Paillusson, S., Mueller, S., Sancho, R.M., Lau, K.F., Vizcay-Barrena, G., Lin, W.L., Xu, Y.F., Lewis, J., et al. (2014). ER-mitochondria associations are regulated by the VAPB-PTPIP51 interaction and are disrupted by ALS/FTD-associated TDP-43. Nature communications 5, 3996.

Sui, X., Arlt, H., Brock, K.P., Lai, Z.W., DiMaio, F., Marks, D.S., Liao, M., Farese, R.V., and Walther, T.C. (2018). Cryo–electron microscopy structure of the lipid droplet–formation protein seipin. J Cell Biol 217, 4080–4091.

Szymanski, K.M., Binns, D., Bartz, R., Grishin, N.V., Li, W.-P., Agarwal, A.K., Garg, A., Anderson, R.G., and Goodman, J.M. (2007). The lipodystrophy protein seipin is found at endoplasmic reticulum lipid droplet junctions and is important for droplet morphology. Proceedings of the National Academy of Sciences 104, 20890–20895.

Thiam, A.R., and Ikonen, E. (2020). Lipid Droplet Nucleation. Trends in Cell Biology.

Thiam, A.R., Farese, R.V., Jr., and Walther, T.C. (2013). The biophysics and cell biology of lipid droplets. Nature reviews Molecular cell biology 14, 775–786.

Wang, H., Becuwe, M., Housden, B.E., Chitraju, C., Porras, A.J., Graham, M.M., Liu, X.N., Thiam, A.R., Savage, D.B., and Agarwal, A.K. (2016). Seipin is required for converting nascent to mature lipid droplets. Elife 5, e16582.

Wang, S., Idrissi, F.-Z., Hermansson, M., Grippa, A., Ejsing, C.S., and Carvalho, P. (2018). Seipin and the membrane-shaping protein Pex30 cooperate in organelle budding from the endoplasmic reticulum. Nature Communications 9, 2939.

Welte, M.A., and Gould, A.P. (2017). Lipid droplet functions beyond energy storage. Biochim Biophys Acta Mol Cell Biol Lipids 1862, 1260–1272.

Windpassinger, C., Auer-Grumbach, M., Irobi, J., Patel, H., Petek, E., Hörl, G., Malli, R., Reed, J.A., Dierick, I., and Verpoorten, N. (2004). Heterozygous missense mutations in BSCL2 are associated with distal hereditary motor neuropathy and Silver syndrome. Nature Genetics 36, 271.

Wolinski, H., Hofbauer, H.F., Hellauer, K., Cristobal-Sarramian, A., Kolb, D., Radulovic, M., Knittelfelder, O.L., Rechberger, G.N., and Kohlwein, S.D. (2015). Seipin is involved in the regulation of phosphatidic acid metabolism at a subdomain of the nuclear envelope in yeast. Biochimica et Biophysica Acta (BBA)-Molecular and Cell Biology of Lipids 1851, 1450–1464.

Yan, R., Qian, H., Lukmantara, I., Gao, M., Du, X., Yan, N., and Yang, H. (2018). Human SEIPIN binds anionic phospholipids. Developmental Cell 47, 248–256.

Zoni, V., Khaddaj, R., Campomanes, P., Thiam, A.R., Schneiter, R., and Vanni, S. (2021). Pre-existing bilayer stresses modulate triglyceride accumulation in the ER versus lipid droplets. Elife 10, e62886.

